# Sulfatase modifying factors control the timing of zebrafish convergence and extension morphogenesis

**DOI:** 10.1101/2025.10.09.681375

**Authors:** Ailen Soledad Cervino, Amrita Basu, Ryan J. Weiss, Gursimran Kaur Bajwa, Rubén Marín Juez, Sandra Grimm, Cristian Coarfa, Margot Kossmann Williams

## Abstract

To shape the emerging body plan, morphogenetic cell movements must be coordinated not only in space, but also in time. Convergence and Extension (C&E) movements that elongate the anteroposterior axis initiate with precise timing during vertebrate gastrulation, but the mechanisms controlling their onset remain unknown. We examined this question using zebrafish embryonic explants that faithfully recapitulate C&E cell movements and their precise timing in culture, in isolation from other gastrulation movements. We determined that new transcription is required at gastrulation onset for C&E in explants and identified *sulfatase modifying factor 2* (*sumf2*) as a candidate ‘trigger’ gene expressed at this time. *sumf2* and its paralog *sumf1* encode negative and positive regulators, respectively, of all sulfatase enzymes, which remove sulfate groups from their substrates, altering their biological activity. In zebrafish embryos and explants, *sumf1 and sumf2* expression levels invert at gastrulation onset, predicting a reduction in sulfatase activity and consequent increase of substrate sulfation. We found that overexpressing *sumf1* and *sumf2* causes delayed or precocious C&E onset, respectively, whereas loss of *sumf1* and *sumf2* function shifts C&E timing in the opposite direction. We further identified Sulf1, an extracellular sulfatase that modifies heparan sulfate proteoglycans (HSPGs), as the key effector by which *sumf1* and *sumf2* control C&E timing. Accordingly, reduced or increased levels of sulfated heparan sulfate similarly shift C&E onset and suppress *sumf1* and *sumf2* mutant phenotypes. Together, our work supports a model in which *sumf2* expression at zebrafish gastrulation onset reduces sulfatase activity, rewriting HSPG sulfation patterns to promote and/or permit C&E morphogenesis.

## INTRODUCTION

Morphogenetic cell movements must be coordinated not only in space, but also in time to properly shape the embryonic body plan. Indeed, changes in the timing of developmental events, termed heterochrony, can underlie malformations in individuals and fuel evolutionary change in populations (*1, 2*). One striking example of morphogenetic timing is the onset of gastrulation cell movements that form the primordial germ layers and shape them into the nascent embryonic axes. In many species, multiple gastrulation movements occur simultaneously and/or in rapid succession, such that the absolute timing of one process is necessary to preserve its timing relative to the others. For example, many teleost fish and amphibian embryos exhibit epiboly (which thins and spreads the epiblast), internalization (which brings mesoderm and endoderm germ layers inside the embryo), and convergence & extension (C&E) (which elongate the anteroposterior axis) movements simultaneously. The onset of epiboly in zebrafish is thought to result from a rapid fluidization of the epiblast, a consequence of cell rounding during meta-synchronous cell cleavages (*3, 4*). Initial internalization of mesoderm and endoderm cells may be triggered by a threshold level of Nodal signaling activity (*5*), which promotes cell protrusions that “un-jam” cells to enable them to ingress at the margin (*6*) or ectopically (*7*). Once leader cells ingress at the margin, “followers” in adjacent spatial domains subsequently internalize in a temporally ordered fashion according to their expression of *hox* genes (*8*), similar to a mechanism reported during chick gastrulation (*9*). However, mechanisms controlling the timing of C&E morphogenesis remain elusive.

C&E movements simultaneously narrow and elongate embryonic tissues via polarized cell rearrangements, providing the major driving force of anteroposterior (AP) axis extension and, in many species, neural tube closure (*10–13*). C&E are driven by polarized cell behaviors including mediolateral (ML) cell elongation and alignment, ML-biased cell protrusions, and polarized contraction of cell interfaces by which cells exchange neighbors to form a longer and narrower array (*14, 15*). Vertebrate C&E movements and/or their underlying ML intercalation behaviors often begin at mid-gastrulation (*16–18*). In zebrafish, this is marked by a switch in the trajectory of cell movements toward the embryo’s dorsal side at ∼75% epiboly (∼8 hours post fertilization (hpf)) (*18, 19*). Evidence suggests that these movements do not result from chemotaxis (*20*), but rather from spatially restricted cell-cell interactions whose domains are defined by morphogen signaling gradients (*21–23*). The signature ML polarity of vertebrate C&E behaviors is under control of planar cell polarity (PCP) signaling, which orients cell behaviors with respect to the embryonic axes (*24, 25*). This results from the polarized membrane localization of core PCP signaling components (*26–29*), which directly precedes the onset of C&E cell behaviors (*22, 30*). The precision and coordination of these events imply the existence of a timing cue that determines the onset of C&E cell behaviors. Several morphogen signaling pathways are essential for C&E, with BMP preventing C&E of the ventral mesoderm (*21, 31, 32*) and Nodal and FGF signaling promoting C&E dorsally (*22, 33–37*). However, all these pathways are active hours before C&E begins at mid-gastrulation, suggesting that their activity might not be directly responsible for the timing of C&E onset. Indeed, although Activin (which signals through the Nodal pathway) was sufficient to induce axial mesoderm exhibiting C&E in *Xenopus* animal cap explants, the timing of C&E remained constant regardless of when Activin was applied (*38*). This suggests that the timing mechanism functions in parallel with known C&E regulators, but its molecular basis is unknown.

Mechanisms controlling the timing of other early embryonic events have been extensively studied, including early cell cycles and zygotic genome activation (ZGA). For example, early cell cleavages are controlled by cyclical expression and degradation of maternally expressed Cyclins (*39*). After a set number of cell divisions, ZGA is triggered by a threshold nuclear/cytoplasmic (N/C) ratio (*40, 41*) (*41–43*), which is thought to function through titration of histones and other nuclear factors (*42–45*). In amphibian embryos, the timing of gastrulation was linked not to fertilization or ZGA, but to the first embryonic cleavage (*46, 47*), and this timing is reportedly regulated predominantly by cytoplasmic factors (*46, 48–50*). However, the onset of gastrulation morphogenesis is unchanged in embryos with varying cell size and cell cycle length (*50–53*), making mechanisms that count cell cycles or measure N/C ratios unlikely regulators of C&E timing. Zebrafish gastrulation morphogenesis requires zygotic transcription (*54*), but its timing is uncoupled from that of ZGA, instead supporting a model by which new gene expression triggers C&E at a specific time.

Using a reductive zebrafish embryonic explant model in which C&E is isolated from the other gastrulation cell movements, we determined that new transcription is required at gastrulation onset for C&E to occur, and that *sulfatase modifying factor 2* (*sumf2*) is initially expressed during this critical window in both explants and intact embryos. *sumf1* and *sumf2* encode Formylglycine Generating Enzyme (FGE) (*55–58*) and its antagonist and paralog pFGE (*59–61*), respectively, a pair whose balance determines the activity of every sulfatase enzyme in the body (*62–65*). *sumf1* is maternally expressed, and its transcript abundance drops just as *sumf2* is expressed at gastrulation onset, inverting *sumf1/sumf2* levels and altering sulfation in the embryo. We show that overexpression of *sumf1* and *sumf2* causes delayed or precocious C&E onset, respectively, in both explants and embryos, whereas loss of *sumf1* and *sumf2* function shifts C&E timing in the opposite direction. We further identified Sulf1 as a key sulfatase by which Sumf1 and Sumf2 modify C&E timing. Reduced or increased levels of sulfated heparan sulfate, the predominant substrate of Sulf1 (*66–68*), similarly shifts C&E onset and suppresses *sumf1* and *sumf2* mutant phenotypes, indicating that heparan sulfate proteoglycans (HSPGs) mediate C&E timing downstream of sulfatase modifiers. Together, these data support a model in which *sumf2* expression at gastrulation onset reduces sulfatase activity, triggering a switch in HSPG sulfation patterns to promote and/or permit C&E morphogenesis.

## RESULTS

### *Ex vivo* convergence & extension requires new transcription at gastrulation onset

Blocking transcription prior to ZGA prevents all gastrulation morphogenesis in zebrafish embryos (*54*). However, it was not determined when new transcription is required for gastrulation cell movements nor the specific genes that are required. We previously showed that otherwise naïve zebrafish embryonic explants expressing Nodal signaling components recapitulate both C&E behaviors and their precise timing in culture, while lacking other gastrulation movements (*36*), making them an ideal system to study C&E timing. To identify the temporal window of gene expression required for C&E, we treated explants expressing the constitutively active Nodal receptor Acvr1b*\** (also known as TARAM-A* (*69*)) with a time-course of the irreversible transcription inhibitor Triptolide (*70*). We then assessed their ability to undergo C&E by measuring their extension when sibling embryos reached the 4-somite stage (12 hpf) (**Fig. 1A**). *acvr1b\** explants treated when intact siblings reached 50% epiboly (5.3 hpf) failed entirely to extend, while those treated shortly thereafter at shield stage (6 hpf, gastrulation onset) or later were able to extend, albeit incompletely (**Fig. 1B, C**). To rule out the possibility that Triptolide interferes with C&E by disrupting Nodal signaling dynamics, we repeated the experiment by activating Nodal under two additional experimental conditions. First, we expressed the Nodal ligand Ndr2/Cyc in explants, which display delayed Nodal activation compared to *acvr1b\** explants (*71*). Second, we expressed *acvr1b\** in explants generated from Nodal signaling-deficient maternal-zygotic (MZ)*oep*-/- embryos (*72*), which exhibit continued signaling in the absence of feed-forward synthesis of new Nodal ligands. Both conditions responded identically to wildtype (WT) *acvr1b\** explants, indicating that the requirement for transcription is independent of Nodal signaling *per se* (**Supplementary Fig. 1A, B, D, E**). This is consistent with Activin-induced *Xenopus* animal cap explants extending on time regardless of when in the competence window Activin was added (*38*). Finally, WT *acvr1b\** explants treated with the reversible transcription inhibitor Flavopiridol (*73*) at 50% epiboly and washed out at 90% epiboly (5.3–9 hpf) similarly failed to extend (**Supplementary Fig. 1C, F**), demonstrating that the stage of inhibition, rather than its duration, was responsible for failed C&E. These results indicate that new gene expression starting at gastrulation onset is required for *ex vivo* C&E movements, independent of Nodal signaling dynamics.

**Fig. 1.**
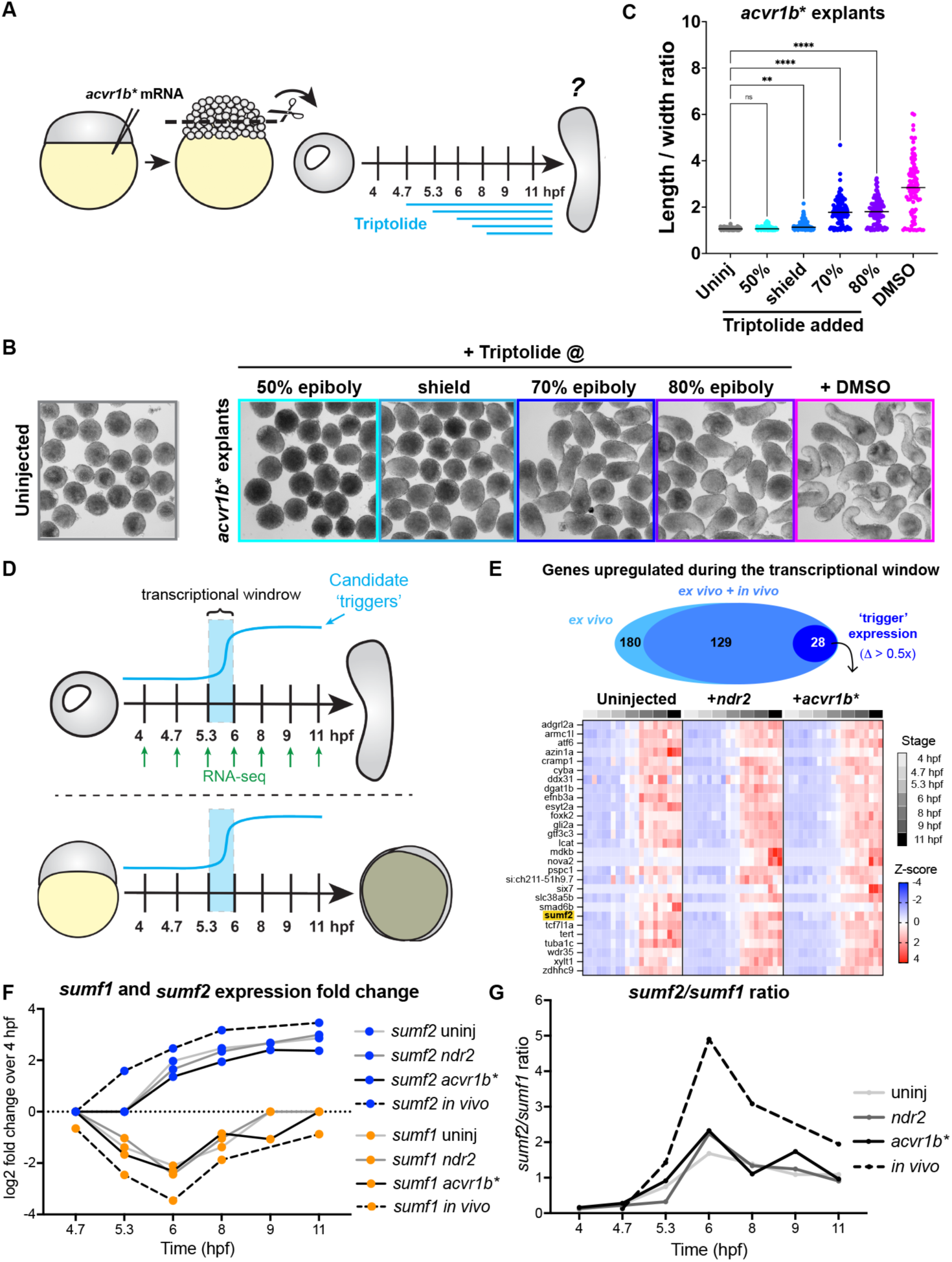
*Ex vivo* convergence & extension requires new transcription at gastrulation onset. **(A)** Diagram of embryonic explants and Triptolide treatments (modified from (*36*)). **(B)** Representative images of uninjected and *acvr1b\** explants at 12 hpf (equivalent of 4-somite stage) after treatment with triptolide at the indicated stages or with DMSO at 50% of epiboly. **(C)** Length/width ratios of explants shown in (B). Each dot represents a single explant from three independent trials, black bars are median values; p<0.0001, Mann-Whitney test. **(D)** Overview of comparisons between published bulk RNA-sequencing experiments from seven developmental stages in three explant conditions (uninjected, *acvr1b⁎*, and *ndr2*) (top) and intact embryos (bottom). Candidate ‘trigger’ genes were strongly upregulated within the previously determined transcriptional window (5.3–6 hpf). **(E)** (Top) 180 genes exhibited ‘trigger’ expression patterns in all explant conditions (light blue); of these, 129 were also upregulated in intact embryos (blue), and 28 showed a sharp increase (Δ > 0.5x) with significant expression levels (>5 TPM, dark blue). (Bottom) Heatmap of candidate ‘trigger’ genes in uninjected*, acvr1b\**, and *ndr2* explants. *sumf2* (highlighted in yellow) was selected for further study. **(F)** Fold-change expression of *sumf1* and *sumf2* transcripts over 4 hpf in explants (solid lines) and embryos (dashed lines) over time. **(G)** *sumf2/sumf1* expression ratio in explants (solid line) and embryos (dashed line) over time. Note the peak in the *sumf2/sumf1* ratio at 6 hpf, coinciding with the previously determined transcriptional window.

**Supplementary Fig. 1.**
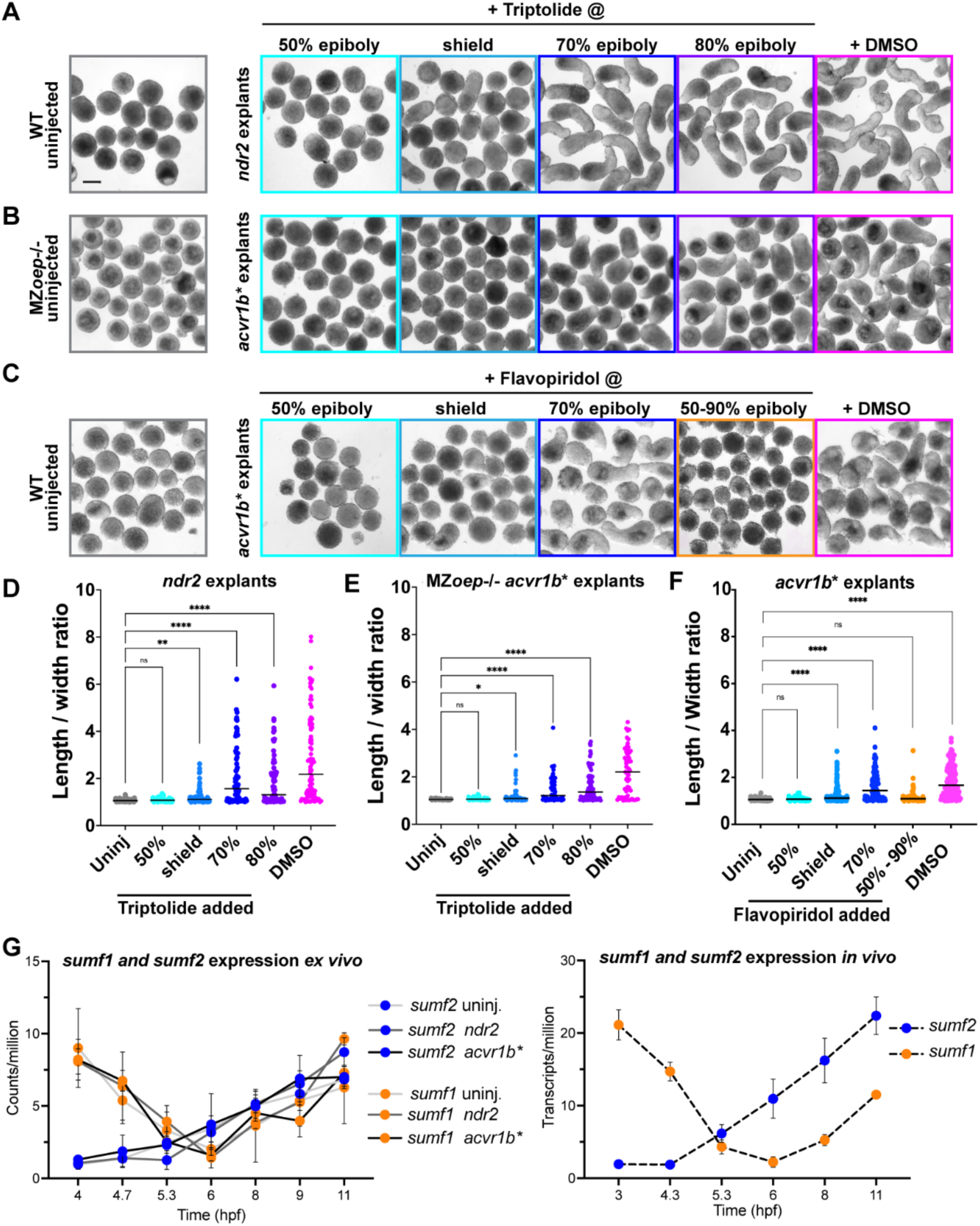
*Ex vivo* convergence & extension requires new gene expression at gastrulation onset independently of Nodal signaling dynamics. **(A, B)** Representative images of WT uninjected and *ndr2* explants (A), or MZ*oep*-/- uninjected and *acvr1b\** explants (B) at 12 hpf (equivalent of 4-somite stage) after treatment with triptolide at the indicated stages or with DMSO at 50% of epiboly. Scale bar 200 μm. **(C)** Representative images of WT uninjected and *acvr1b\** explants at 12 hpf (equivalent of 4-somite stage) after treatment with reversible transcription inhibitor Flavopiridol at the indicated stages or with DMSO at 50% of epiboly. **(D-F)** Length/width ratios of explants shown in (A, B and C), respectively. Each dot represents a single explant from three independent trials, black bars are median values; p<0.0001, Mann-Whitney test. **(G)** Expression levels of *sumf1* and *sumf2* in uninjected (light grey lines), *ndr2* (grey lines) and *acvr1b\** (black lines) explants (left), and embryos (dashed lines, right) over time. Dots represent the mean of three (for explants) or five (for embryos) bulk RNA-seq replicates and error bars represent standard deviation.

### *sumf1* and *sumf2* transcript levels invert at gastrulation onset

Having established that transcription at gastrulation onset is crucial for C&E, we next identified genes whose expression is strongly upregulated during this time window. We previously profiled transcription in *ndr2, acvr1b*,* and uninjected control explants by bulk RNA sequencing at seven developmental stages spanning gastrulation (*71*) (**Fig. 1D**). We examined these data for candidate “trigger” genes whose expression exhibited a substantial increase between 50% epiboly and shield stage, corresponding to our experimentally determined transcriptional window. Because the timing of C&E onset is independent of Nodal signaling dynamics, we only considered genes with similar temporal expression profiles in explants of all three conditions, including uninjected controls. This analysis yielded a list of 180 genes, 129 of which had similar increasing expression profiles in intact zebrafish gastrulae (*74*) (**Supplementary Table 1A**). From these, we selected genes exhibiting at least a 50% increase (Δ > 0.5x) in expression at gastrulation onset (shield stage) from their baseline at 50% epiboly and whose expression at shield stage *in vivo* was over 5 transcripts per million, producing a list of 28 genes (**Fig. 1E**) (**Supplementary Table 1B**). Because C&E begins simultaneously in intact embryos and explants, we hypothesized these 28 genes represent candidates whose expression at gastrulation onset may trigger C&E cell behaviors.

Among our candidate genes was *sulfatase modifying factor 2* (*sumf2*). *sumf2* encodes pFGE, a paralog and antagonist of Formylglycine Generating Enzyme (FGE) (*65, 75*), encoded by *sulfatase modifying factor 1* (*sumf1*). As its name implies, FGE converts cysteine residues to formylglycine, a rare but essential post-translational modification required within the active site of all sulfatase enzymes for their activity (*75*). Sulfatases catalyze the removal of sulfate groups from a variety of substrates, including lipids, steroids, and glycosaminoglycans, modifying their biological activity (*76, 77*). Both pFGE (*sumf2*) and FGE (*sumf1*) are ER-resident proteins that share high amino acid sequence similarity with the main distinction that pFGE lacks enzymatic activity (*55, 75*). Evidence suggests that pFGE binds FGE alone or in complex with its sulfatase substrates (*59, 65*), but it is not clear precisely how pFGE (*sumf2*) antagonizes FGE (*sumf1*) function. Although other candidates exhibited more dramatic expression increases at gastrulation onset, we selected *sumf2* for further study because its function during vertebrate development has never been explored, and its intriguing complementary temporal expression patterns with *sumf1* during gastrulation. In zebrafish embryos and explants, *sumf1* is maternally expressed but exhibits a sharp reduction in transcript levels just prior to gastrulation onset, coinciding with increased *sumf2* (*74*) (**Fig. 1F, Supplementary Fig. 1G**). Because *sumf2* and *sumf1* encode a negative and positive regulator of sulfatase activity, respectively, we hypothesized that this peak in the *sumf2/sumf1* ratio at gastrulation onset (6hpf) (**Fig. 1G**) reduces sulfatase activity and thus increases sulfation of their substrates to trigger C&E movements.

### Excess or loss of sulfatase modifying factors causes C&E defects in zebrafish gastrulae

To investigate the role of sulfatase modifying factors in C&E morphogenesis, we employed both gain- and loss-of-function approaches and performed morphometric analysis at the end of gastrulation (tailbud stage, 10 hpf). We first overexpressed *sumf1* or *sumf2* by mRNA injection into single-cell WT embryos. *sumf1* overexpressing (OE) gastrulae exhibited a significant reduction in anteroposterior (AP) axis length and wider notochords, characteristic of C&E defects, and *sumf2* OE embryos showed similar but milder phenotypes (**Fig. 2**). Next, we examined gastrulae of the recently characterized *sumf1^la015919Tg^*mutant line (hereafter *sumf1*-/-) harboring a viral insertion in exon 1 that abolishes FGE activity (*78*). Homozygous mutant embryos survived to adulthood (*78*), enabling examination of MZ*sumf1*-/- gastrulae, which also exhibited reduced AP axis length and wider notochords indicative of impaired C&E (**Fig. 2**). Because no existing *sumf2* mutant line was available, we used CRISPR to generate a 7 bp insertion in exon 2 of *sumf2* (named *bcm126*), resulting in a premature stop codon (**Supplementary Fig. 2A**). Homozygous *sumf2^bcm126/bcm126^* embryos (hereafter *sumf2*-/-), showed a substantial reduction in *sumf2* transcript levels by qRT-PCR (**Supplementary Fig. 2C**), confirming this as a strong loss-of-function allele. These mutants also survived to adulthood, enabling us to maintain the line as homozygotes and analyze MZ*sumf2*-/- embryos. Notably, approximately 10% of MZ*sumf2*-/- larvae displayed a tail-curled-up phenotype at 48 hpf and a variable number of *sumf2*-/- adults were undersized and scoliotic (**Supplementary Fig. 2D, E**). Although MZ*sumf2*-/- mutant gastrulae did not exhibit C&E defects, *sumf1* OE in this background produced more severe C&E phenotypes than in WT embryos (**Figure 2**), supporting an antagonistic role for *sumf2*/pFGE on *sumf1*/FGE. Together, these results demonstrate that altering levels of *sumf1* or/and *sumf2* disrupts C&E gastrulation morphogenesis *in vivo*.

**Fig. 2.**
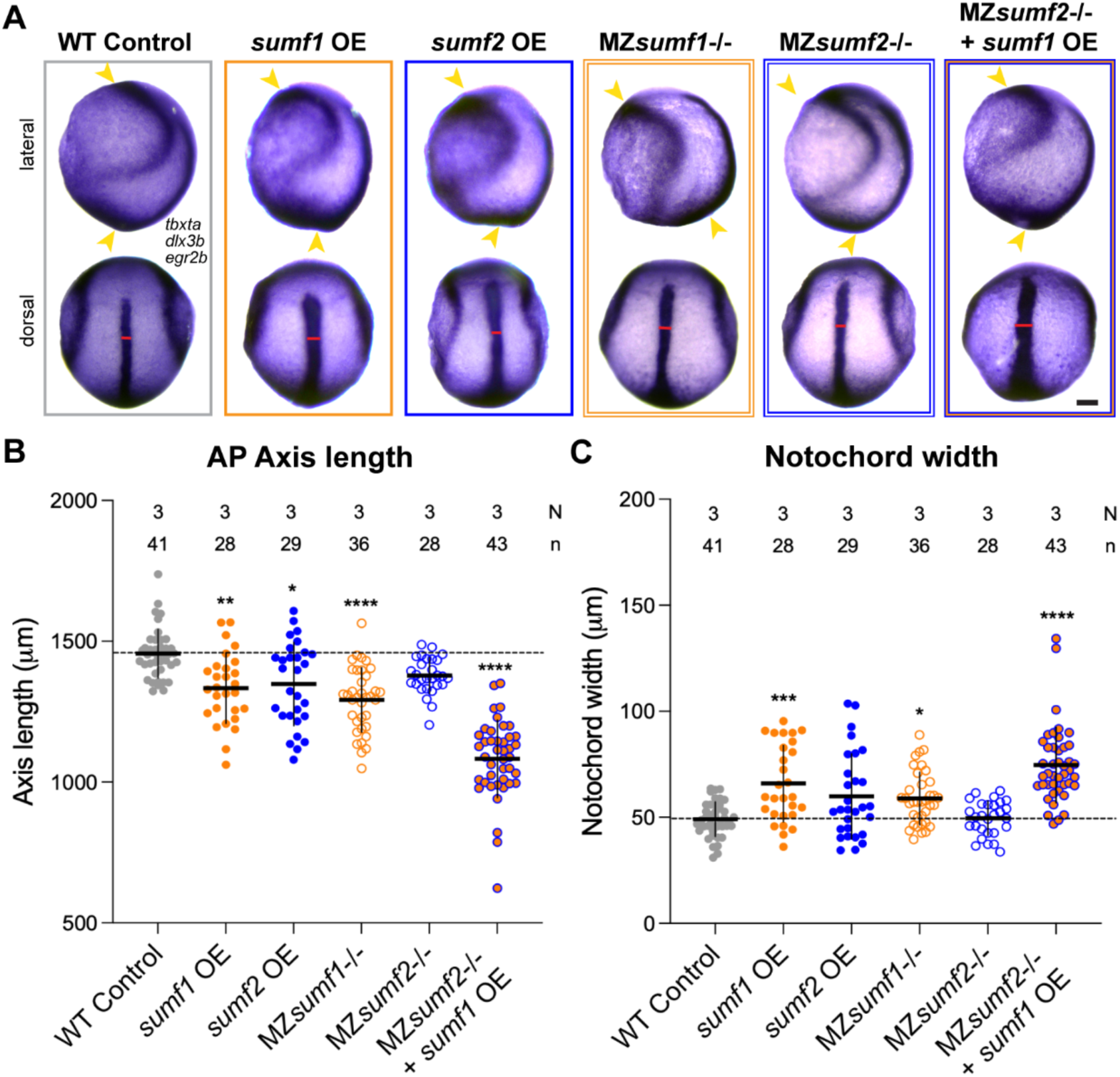
C&E defects in zebrafish gastrulae with altered sulfatases modifiers factor levels. **(A)** Representative images of whole mount in situ hybridization (WISH) for *tbxta* (mesoderm), *dlx3b* (neural plate border) and *egr2b* (rhombomeres 3 & 5) in tailbud stage (10 hpf) embryos of the indicated conditions. Anterior is up in all images, lateral views are shown on top, dorsal views on bottom. Scale bar 100 μm. **(B, C)** Anteroposterior (AP) axis length (B, yellow arrowheads) and notochord width (C, red lines) of embryos depicted in (A). Each dot represents a single embryo, N: number of independent experiments, n: number of embryos. Means and standard deviation are indicated; * p<0.05, ** p<0.01, **** p<0.0001 compared to WT control group by Kruskal–Wallis and Dunn’s multiple comparisons tests.

**Supplementary Fig. 2.**
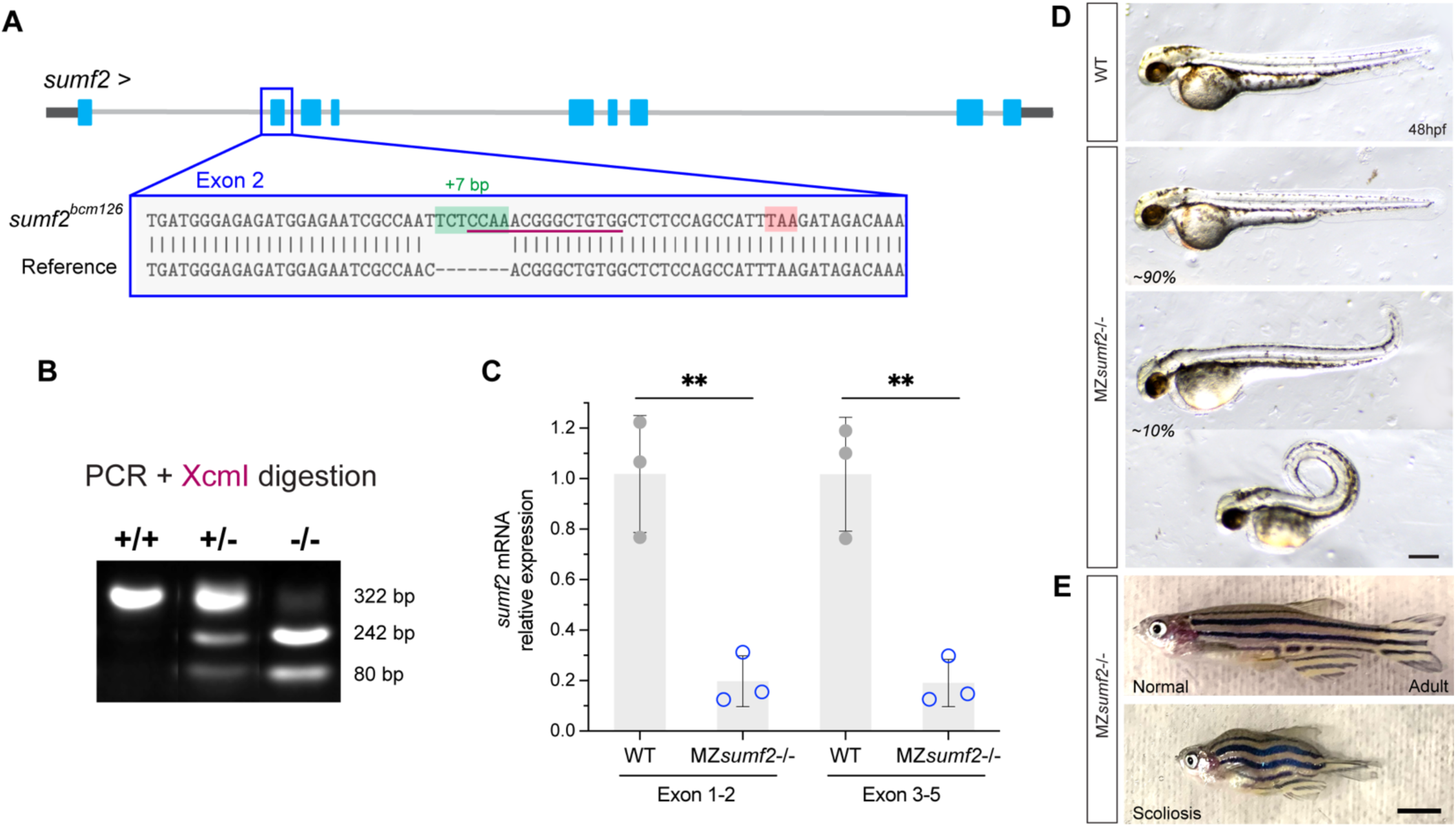
Characterization and genotyping of new *sumf2* loss-of-function allele. **(A)** Diagram of CRISPR-induced 7 bp insertion (green) in exon 2 of *sumf2* (*sumf2^bcm126^)*, resulting in a premature stop codon (red). **(B)** The *sumf2^bcm126^* allele creates an XcmI restriction site (purple underline in (A)), enabling genotyping by digestion of a 322 bp PCR-amplified fragment, which yields 242 bp and 80 bp products in heterozygous (+/−), and homozygous (−/−) but not WT (+/+) samples. **(C)** Relative expression of *sumf2* transcripts in WT and MZ*sumf2*-/- larvae, measured by RT-qPCR using primers spanning *sumf2* exon 1–2 and exon 3–5. Means and standard deviation are indicated. Each dot represents one of three independent clutches; ** p<0.01, Mann-Whitney test. **(D)** Representative images of WT and MZ*sumf2*-/- larvae at 48 hpf. Approximately 90% of MZ*sumf2*-/- larvae developed normally, while ∼10% displayed varying degrees of tail-curled-up phenotypes. Anterior is to the left; scale bar 200 μm. **(E)** Representative images of MZ*sumf2*-/- adult fish showing *Normal* and *Scoliosis* phenotypes. Scale bar 500 μm.

### *sumf1* and *sum2* levels modify the timing of C&E *ex vivo*

Next, we examined the effects of altered *sumf1* and *sumf2* levels in zebrafish embryonic explants, in which C&E is isolated from other concurrent morphogenetic processes. Surprisingly, neither OE nor deficiency of *sumf1* or *sumf2* impaired the extension of *acvr1b\** explants at the equivalent of 4-somite stage (**Supplementary Fig. 3**), indicating that they are dispensable for C&E *per se*. To assess whether *sumf1* or *sumf2* levels instead influenced the timing of C&E morphogenesis, we acquired time-lapse recordings of *acvr1b\** explants to precisely determine the onset of morphological extension. Extension onset was defined as the time point when a visible tip first emerged from an initially round explant (**Fig. 3A**). Importantly, all explants were staged relative to blastopore closure in age-matched intact embryos of the same genetic background mounted in the same dish. As previously described (*36*), WT *acvr1b⁎* explants (WT control) and controls co-injected with *superfold-GFP* mRNA (*sf-GFP* OE) initiated extension around 8hpf. Strikingly, extension began significantly later upon *sumf1* OE and significantly earlier upon *sumf2* OE (**Fig. 3B**). Co-overexpression of *sumf1* and *sumf2* restored the normal time of extension onset (**Fig. 3B**), demonstrating that the balance between these factors (rather than their absolute levels) determines the time of C&E onset. In the converse loss-of-function experiments, we found that extension onset was precocious in MZ*sumf1*-/- explants and delayed in MZ*sumf2*-/- explants (**Fig. 3A, B**), consistent with - but opposite to - *sumf1* and *sumf2* OE phenotypes. MZ*sumf1*-/-, MZ*sumf2*-/- double mutant explants phenocopied the precocious extension of MZ*sumf1*-/- single mutants, consistent with *sumf2/*pFGE modulating C&E timing via its role as an antagonist of *sumf1*/FGE. Together, these results demonstrate that the ratio of *sumf1/sumf2* governs the timing of explant extension, with higher *sumf1* delaying and higher *sumf2* advancing C&E onset.

**Fig. 3.**
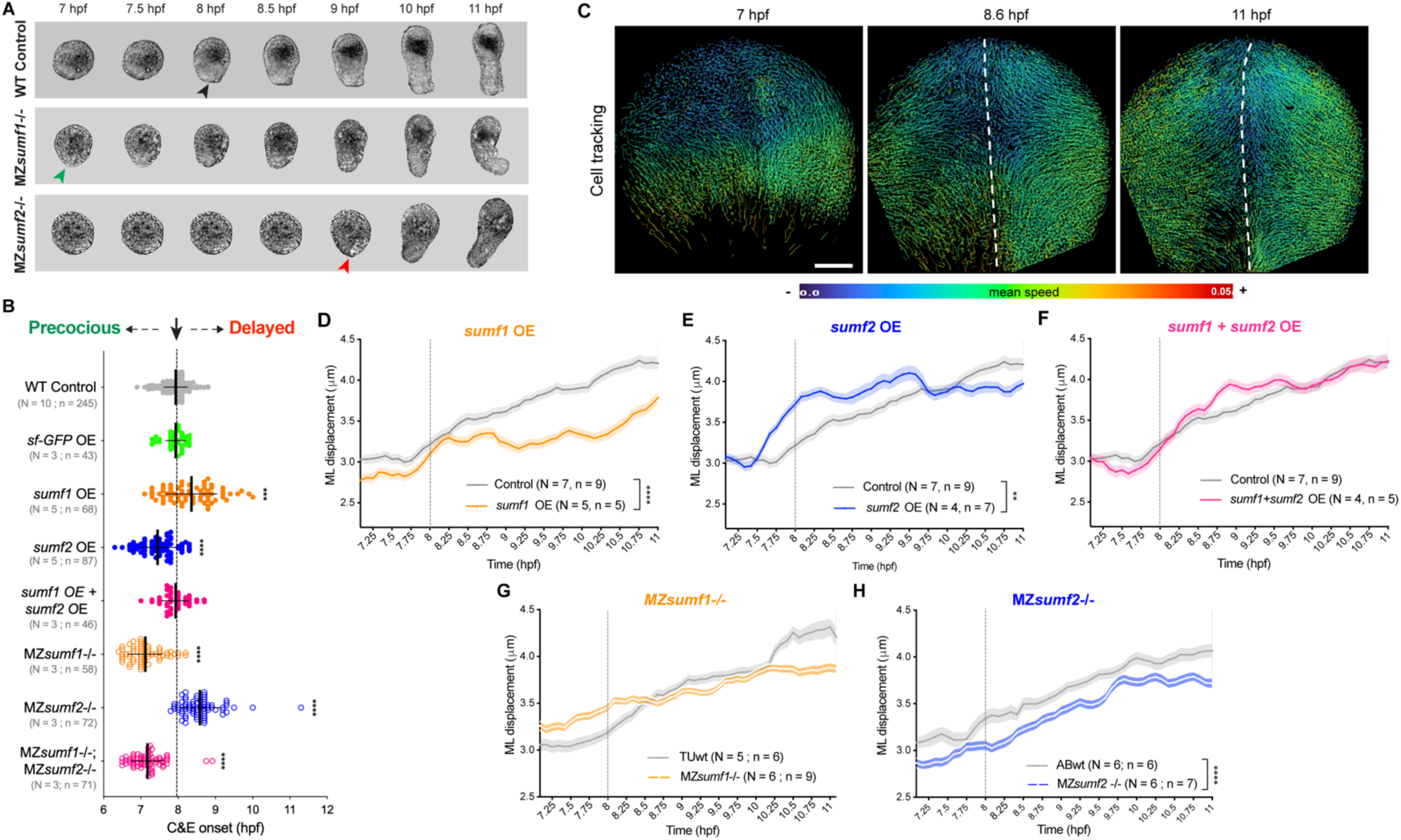
*sumf1* and *sum2* levels control the timing of C&E. **(A)** Representative bright-field images of *acvr1b\** explants of the indicated genotype over time. Black, red, and green arrowheads indicate timely, precocious, and delayed onset of extension, respectively. **(B)** Onset of extension in *acvr1b\** explants of the indicated conditions. Dotted line shows typical extension onset of WT control explants around 8 hpf. Each dot represents a single explant. Means and standard deviation are indicated, N: number of independent experiments, n: number of explants, *** p<0.001, **** p<0.0001 compared with WT control group by Kruskal–Wallis and Dunn’s multiple comparisons tests. **(C)** Representative images of automated nuclear tracking in the dorsal hemisphere of zebrafish gastrulae, starting before C&E onset (7 hpf). Tracks are color-coded by mean speed; dashed lines mark the dorsal midline; scale bar 100 μm. **(D-H)** ML cell displacement over time for WT control (gray lines) and *sumf1* OE (D), *sumf2* OE (E), *sumf1 + sumf2* OE (F), MZ*sumf1*-/- (G), and MZ*sumf2*-/- (H) embryos. Dotted line shows typical onset of convergence movements in WT control embryos around 8 hpf. Means and standard error are indicated, N: number of independent experiments, n: number of embryos. ** p<0.01, **** p<0.0001 compared with WT control group by Wilcoxon signed-rank test.

**Supplementary Fig. 3.**
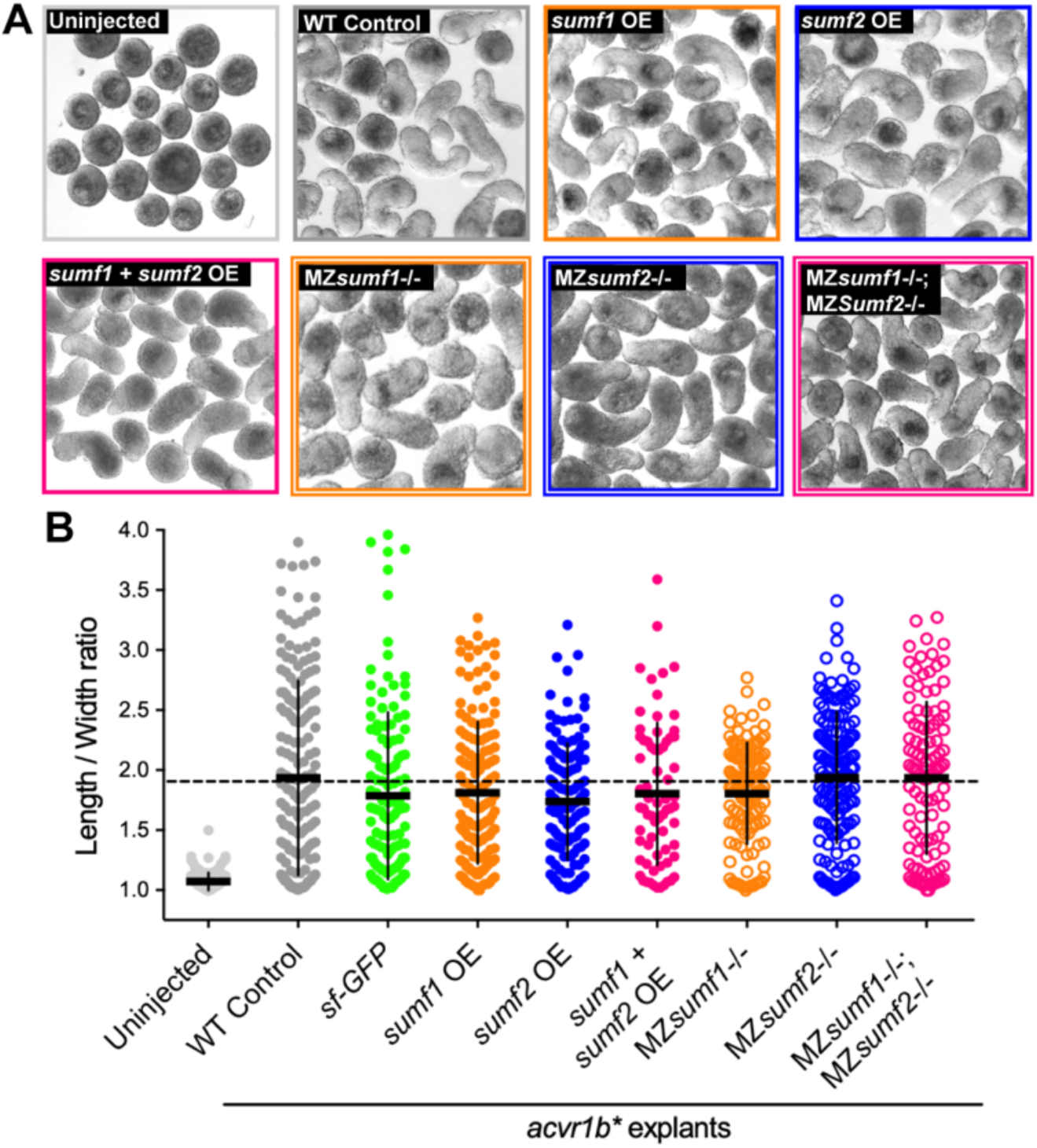
Altered *sumf1* and *sumf2* levels do not impair extension of Nodal-induced explants. **(A)** Representative images of uninjected and *acvr1b\** explants from the indicated experimental conditions. **(B)** Length/width ratios of explants at the equivalent of 4-somite stage (12 hpf) shown in (A). Each dot represents a single explant from three independent trials. Means and standard deviation are indicated. p>0.05, Kruskal–Wallis test.

### *sumf1* and *sum2* levels modify the timing of C&E *in vivo*

To determine whether *sumf1* and *sumf2* similarly modify the timing of C&E cell movements *in vivo*, we acquired confocal time-lapse movies of *H2B-scarlet*-labeled gastrulae from 6.5 – 11 hpf and performed automated nuclear tracking (see *Methods*; **Fig. 3C**). To ensure accurate stage matching across experiments, all embryos were staged relative to formation of the second somite, which was readily visible at the end of each movie. Convergence movements were quantified as the mediolateral (ML) displacement of dorsal and lateral cells over time (**Fig. 3D–H**). Because convergence movements are reduced within the embryonic midline where extension is more prominent (*79*), cells within 100 μm of the dorsal midline were excluded from our analysis. In WT control embryos, we observed an upward inflection of ML displacement beginning around 7.75 - 8 hpf, marking the onset of convergence movements (consistent with previous findings (*18, 19*)), which continued to increase gradually thereafter (**Fig. 3D–H**). Upon *sumf1* OE, baseline convergence speed was reduced and, although convergence movements began at the same time as controls, they remained substantially slower (**Fig. 3D)**. *sumf2* OE embryos, on the other hand, had similar baseline convergence speeds as controls but with an earlier inflection point (**Fig. 3E)**. For much of gastrulation, these movements were also faster than those of controls, indicating a likely change in both onset and pace of convergence movements. As in explants, co-OE of both *sumf1* and *sumf2* restored convergence movements to control timing and speeds (**Fig. 3F)**. Also as in explants, *in vivo* cell tracking of MZ*sumf1-/-* and MZ*sumf2-/-* mutants revealed opposite effects to their overexpression. Convergence onset in MZ*sumf1*-/- embryos trended earlier than controls and baseline speed trended higher, although these movements plateaued later in gastrulation (**Figure 3G)**. Conversely, baseline convergence in MZ*sumf2-/-* mutants was slower than controls with a later inflection point, indicating delayed convergence (**Fig. 3H)**. Together with our *ex vivo* results, these data support a model in which *sumf2*/pFGE counters the activity of *sumf1*/FGE to control the onset and pace of C&E cell movements during gastrulation.

### Sulfatase modifiers govern C&E via the extracellular sulfatase Sulf1

Because *sumf1*/FGE and *sumf2*/pFGE together regulate sulfatase activity levels, we hypothesized that high *sumf2/sumf1* ratios trigger C&E onset by reducing the activity of one or more key sulfatases. Indeed, increased sulfatase activity was shown to disrupt gastrulation morphogenesis in *Xenopus* and sea urchin (*64, 80–83*). The zebrafish genome encodes 17 sulfatases, 11 of which are expressed during peri-gastrulation stages (*74*) (**Supplementary Fig. 4A**). To identify which of these mediates the effect of *sumf1/sumf2* on C&E timing, we overexpressed each of these 11 sulfatases and performed morphometric analysis at tailbud stage and scored axis phenotypes at 24 hpf. Notably, three of them — the heparan sulfate endosulfatases *sulf1* and *sulf2a*, and the chondroitin sulfatase *arsb* — produced severe AP axis extension defects and increased notochord width, consistent with C&E defects (**Supplementary Fig. 4B-E**). Unlike *sumf1* and *sumf2* OE, overexpression of these three sulfatases also prevented full extension of *acvr1b⁎* explants at the equivalent of 4-somite stage (**Supplementary Fig. 4F, G**). However, only *sulf1* OE significantly delayed *acvr1b\** explant extension, while the other two had no effect on C&E timing (**Fig. 4A**). Thus, of the 17 zebrafish sulfatases, only Sulf1 was found to affect both C&E morphogenesis and its timing.

**Fig. 4.**
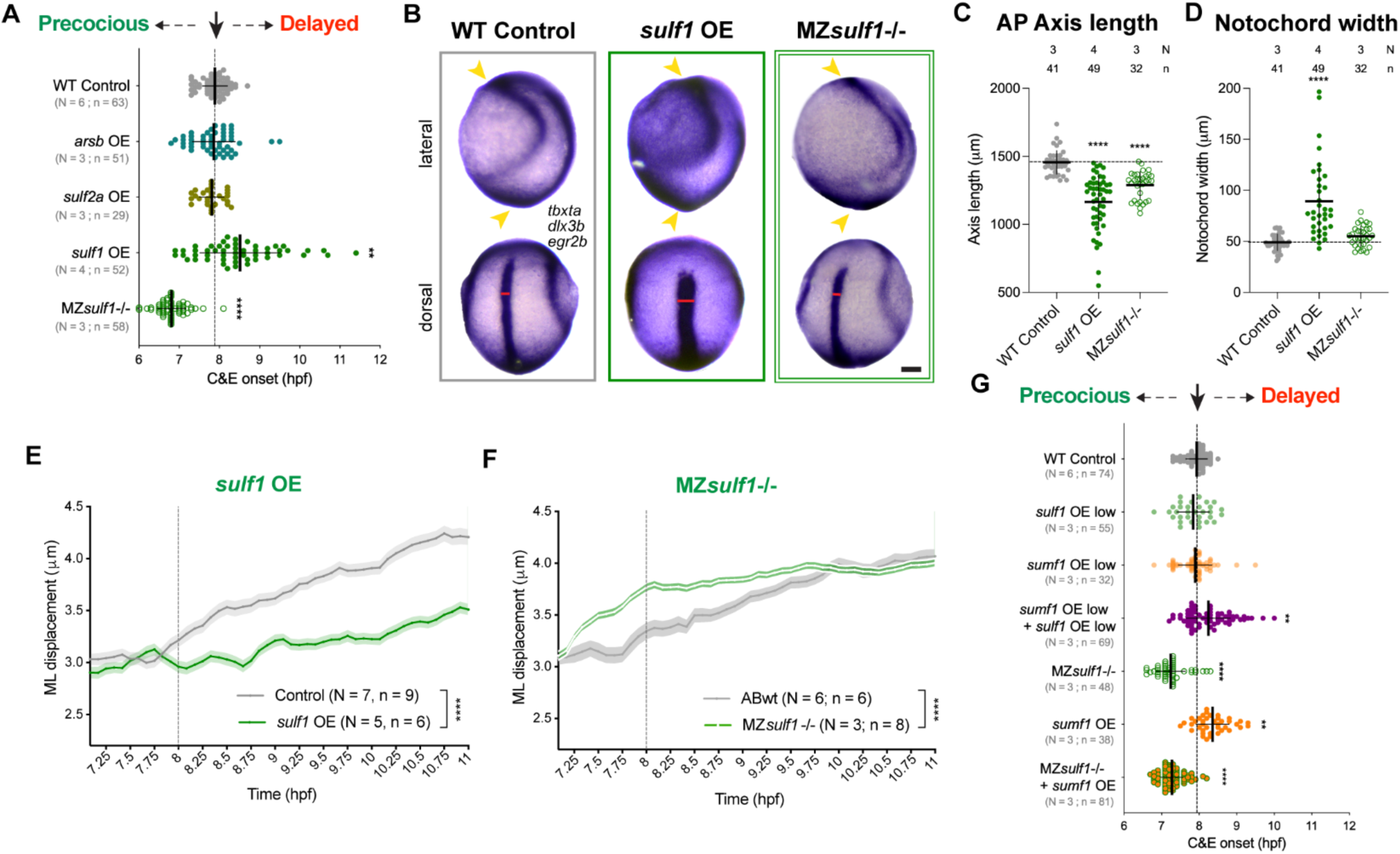
The extracellular sulfatase Sulf1 governs C&E and its timing. **(A)** Onset of extension in *acvr1b\** explants of the indicated conditions. Dotted line shows typical extension onset of WT control explants around 8 hpf. Each dot represents a single explant. Means and standard deviation are indicated, N: number of independent experiments, n: number of explants. *** p<0.001, **** p<0.0001 compared with WT control group by Kruskal–Wallis and Dunn’s multiple comparisons tests. **(B)** Representative images of WISH for *tbxta* (mesoderm), *dlx3b* (neural plate border) and *egr2b* (rhombomeres 3 & 5) in tailbud stage (10 hpf) embryos of the indicated conditions. Anterior is up in all images, lateral views are on top, dorsal views on bottom. Scale bar 100 μm. **(C, D)** Anteroposterior (AP) axis length (C) and Notochord width (D) were quantified as in Figure 2 from embryos depicted in (B). Each dot represents a single embryo. Means and standard deviation are indicated. **** p<0.0001 compared with WT control group by Kruskal–Wallis and Dunn’s multiple comparisons tests. **(E, F)** ML cell displacement from automated nuclear tracking (as in Figure 3) in WT control, *sulf1* OE (E) and MZ*sulf1*-/- (F) zebrafish gastrulae. Means and standard error are indicated. N: number of independent experiments, n: number of embryos. **** p<0.0001 compared with WT control group by Wilcoxon signed-rank test. **(G)** Onset of extension in *acvr1b\** explants of the indicated conditions. Dotted line shows typical extension onset of WT control explants around 8 hpf. Each dot represents a single explant. Means and standard deviation are indicated, N: number of independent experiments, n: number of explants. ** p<0.01, **** p<0.0001 compared with WT control group by Kruskal–Wallis and Dunn’s multiple comparisons tests.

**Supplementary Fig. 4.**
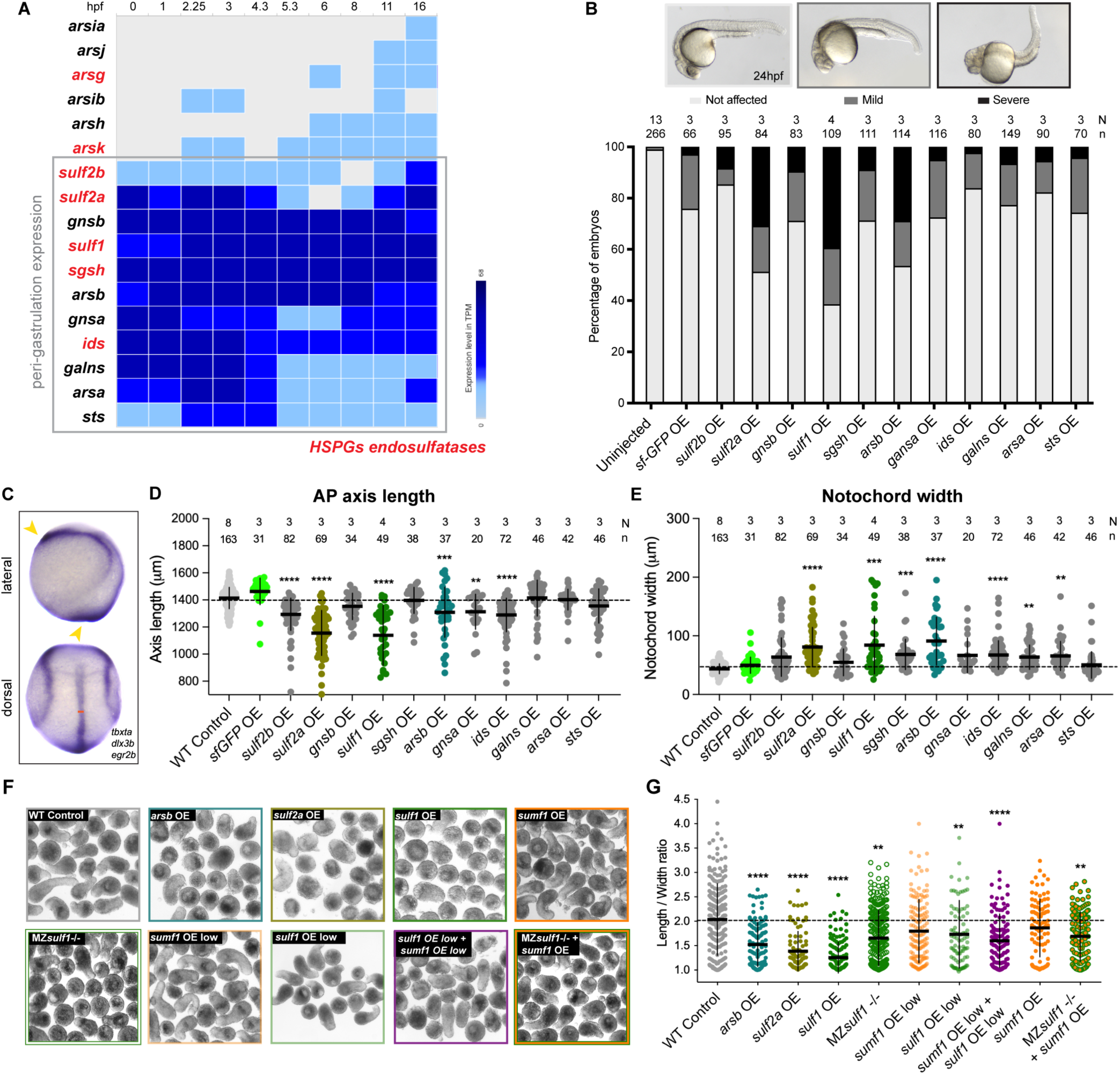
Evaluation of C&E defects upon sulfatase overexpression. **(A)** Expression levels of all 17 zebrafish sulfatases over time based on publicly available bulk RNA-seq data. Gray square indicates the 11 expressed during peri-gastrulation, HSPG endosulfatases are in red. **(B)** (Top) Representative images of 24 hpf embryos overexpressing each of the 11 peri-gastrulation sulfatases displaying a range of axis phenotypes scored as *“Not affected”*, *“Mild”* or *“Severe”*. (Bottom) Mean percentage of 24 hpf embryos of the conditions indicated exhibiting each class of axis defect. N: number of independent experiments, n: number of embryos. **(C)** WISH for *tbxta* (mesoderm), *dlx3b* (neural border) and *egr2b* (rhombomeres 3 & 5) in tailbud stage (10 hpf) embryos shown from lateral (top) and dorsal (bottom) views. **(D, E)** Anteroposterior (AP) axis length (D) and Notochord width (E) (as in Figure 2) in control and sulfatase-overexpressing embryos. Each dot represents a single embryo. Means and standard deviation are indicated. ** p<0.01, *** p<0.001, **** p<0.0001 as compared with WT control group by Kruskal–Wallis and Dunn’s multiple comparisons tests. **(F)** Representative images of *acvr1b\** explants from the indicated experimental conditions. **(G)** Length/width ratios of 12 hpf explants shown in (F). Each dot represents a single explant. Means and standard deviation are indicated. * p>0.05, ** p<0.01, **** p<0.0001 compared with WT control group by Kruskal–Wallis and Dunn’s multiple comparisons tests.

To further investigate the role of Sulf1 in zebrafish gastrulation morphogenesis, we examined C&E in MZ*sulf1^sjr9/sjr9^*full-locus deletion mutants (Kaur Bajwa et al, in submission) (hereafter MZ*sulf1-/-*). MZs*ulf1-/-* embryos exhibited reduced AP axis length at tailbud stage (**Fig. 4B-D)** and MZs*ulf1-/- acvr1b*\* explants showed impaired extension (**Supplementary Fig. 4F-G)**, indicating that both gain and loss of Sulf1 reduces C&E. Strikingly, MZ*sulf1*-/- *acvr1b*\* explants exhibited precocious extension (**Fig. 4A**), similar to MZ*sumf1*-/- and *sumf2* OE explants (but opposite to *sulf1* OE). We next quantified the effect of Sulf1 levels on C&E onset *in vivo*. Convergence speed was drastically reduced upon *sulf1* OE (**Fig. 4E**) while MZ*sulf1*-/- gastrulae exhibited earlier and enhanced convergence movements compared to WT controls (**Fig. 4F**). These results highlight Sulf1 as a strong candidate for the key sulfatase controlling C&E onset.

To examine the relationship between Sulf1 and *sumf1*/FGE, we generated *acvr1b\** explants co-injected with low doses of each mRNA alone or in combination. The combination of otherwise sub-phenotypic doses of *sumf1* and *sulf1* synergistically delayed and reduced explant extension (**Fig. 4G, Supplementary Fig. 4F, G),** consistent with *sumf1*/FGE enhancing Sulf1 activity. Finally, we tested whether *sumf1* OE could delay explant extension in the absence of *sulf1*. We found that although *sumf1* OE significantly delayed extension onset in WT explants, it had no effect on the onset of extension in MZ*sulf1*-/- explants, which remained precocious (**Fig. 4G)**. Together, these results implicate Sulf1 as the primary sulfatase through which *sumf1*/FGE and *sumf2*/pFGE govern the timing of C&E morphogenesis.

### Sulfated heparan sulfate proteoglycans increase with C&E onset

Sulf1 is an extracellular sulfatase that removes 6-*O* sulfation from GlcA-GlcNS6S (D0S6) and IdoA2S-GlcNS6S (D2S6) disaccharide units of heparan sulfate proteoglycans (HSPGs)(*67*). We would therefore expect the inversion of *sumf1*/FGE and *sumf2*/pFGE levels at the beginning of gastrulation to decrease Sulf1 activity, leading to increased sulfated HSPG levels at late gastrulation. Indeed, Alcian Blue staining (at pH ∼1) (*84*) for overall glycosaminoglycan (GAG) sulfation increased dramatically from early (50% epiboly) to late (90% epiboly) gastrulation stages, indicating increased embryo-wide GAG sulfation. Treatment with sodium chlorate, which prevents the formation of the sulfate donor 3’-phosphoadenosine 5’-phosphosulfate (PAPS) (*85, 86*), drastically diminished Alcian Blue staining (**Fig. 5A**), confirming its specificity.

**Fig. 5.**
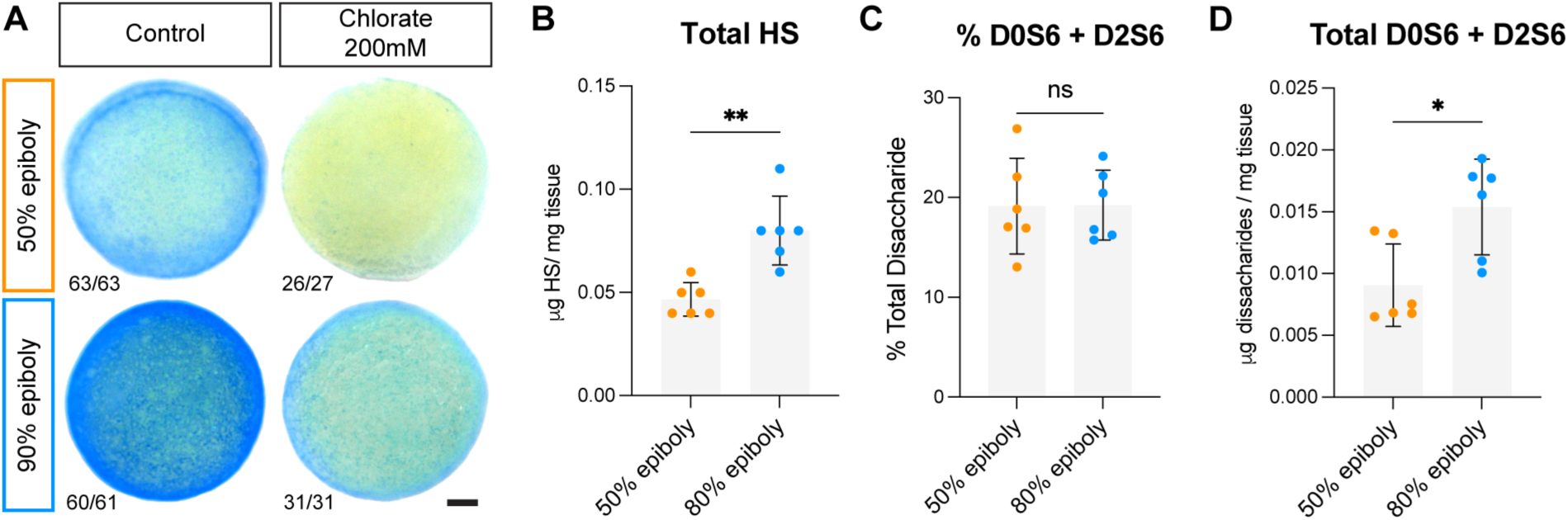
Heparan sulfate proteoglycans and their sulfation increase during gastrulation. **(A)** Representative images of zebrafish early (50% epiboly, 5.3 hpf) and late (90% epiboly, 9 hpf) gastrulae stained with Alcian Blue for sulfated GAGs. Alcian Blue staining was diminished in zebrafish embryos treated at dome stage (4.3 hpf) with 200 mM sodium chlorate. Fractions indicate the number of embryos with the depicted staining pattern over number of embryos examined. Scale bar 100 μm. **(B-D)** HILIC-LC/MS analysis of early and late zebrafish gastrulae showing the total amount of heparan sulfate (HS) (B), the percentage of the D0S6 and D2S6 disaccharides (C), and the total amount of D0S6 and D2S6 disaccharides (D) at the stages indicated. Plots show mean and standard deviation, with each dot representing an independent clutch of 50 embryos, ** p<0.01, * p< 0.05, Mann-Whitney test.

Because Alcian Blue stains all negatively charged GAGs, we next quantified levels of sulfated heparan sulfate (HS) disaccharides by hydrophilic interaction liquid chromatography with mass spectrometry (HILIC-LC/MS) in early and late zebrafish gastrulae (*87*). We found that total HS levels increased significantly between 50% and 80% epiboly stages (**Fig. 5B**). Although we observed no changes in the percentage of the 6-*O* sulfated D0S6 and D2S6 disaccharides between stages (**Fig. 5C**), we did find a significant increase in total levels of these 6-*O* sulfated disaccharides at late gastrulation (reflecting the observed increase in total HS levels) (**Fig. 5D**). These results indicate that sulfation of Sulf1 substrates increases during gastrulation, consistent with our hypothesis that the peak *sumf2/sumf1* ratio at gastrulation onset leads to reduced Sulf1 activity.

### HSPGs mediate the effects of sulfatase modifiers on C&E timing

Sulfatases modify a large number of biological substrates, including lipids, steroids, and multiple GAGs (*88*). To test whether sulfated HSPGs influence C&E morphogenesis and its timing, we experimentally decreased and increased sulfated HS levels within embryos and explants. To inhibit overall sulfation, including GAGs, we treated dome stage embryos (4.3 hpf) with two different doses of sodium chlorate (200 mM and 50 mM), which we previously showed reduced Alcian Blue staining in zebrafish gastrulae (**Fig. 5A**). We observed severe AP axis shortening and cyclopia in 24 hpf embryos treated with the higher dose, and milder axis defects at the lower dose (**Fig. 6A**). A dose-dependent effect on C&E was also observed at tailbud stage (**Fig. 6B-D**), demonstrating that sulfation is required for proper C&E morphogenesis (as previously reported (*89–91*)). Although explants treated with the higher dose were not viable, treatment with the lower dose of sodium chlorate also dramatically reduced extension of *acvr1b⁎* explants (**Supplementary Fig. 5A, B**). Sodium chlorate treatment also significantly delayed extension onset in both WT control and, importantly, in MZ*sumf1*-/- explants in which extension otherwise occurred early (**Fig. 6E**, **Supplementary Fig. 5C**). This finding strongly suggests that loss of *sumf1* induces precocious C&E by enhancing sulfation.

**Fig. 6.**
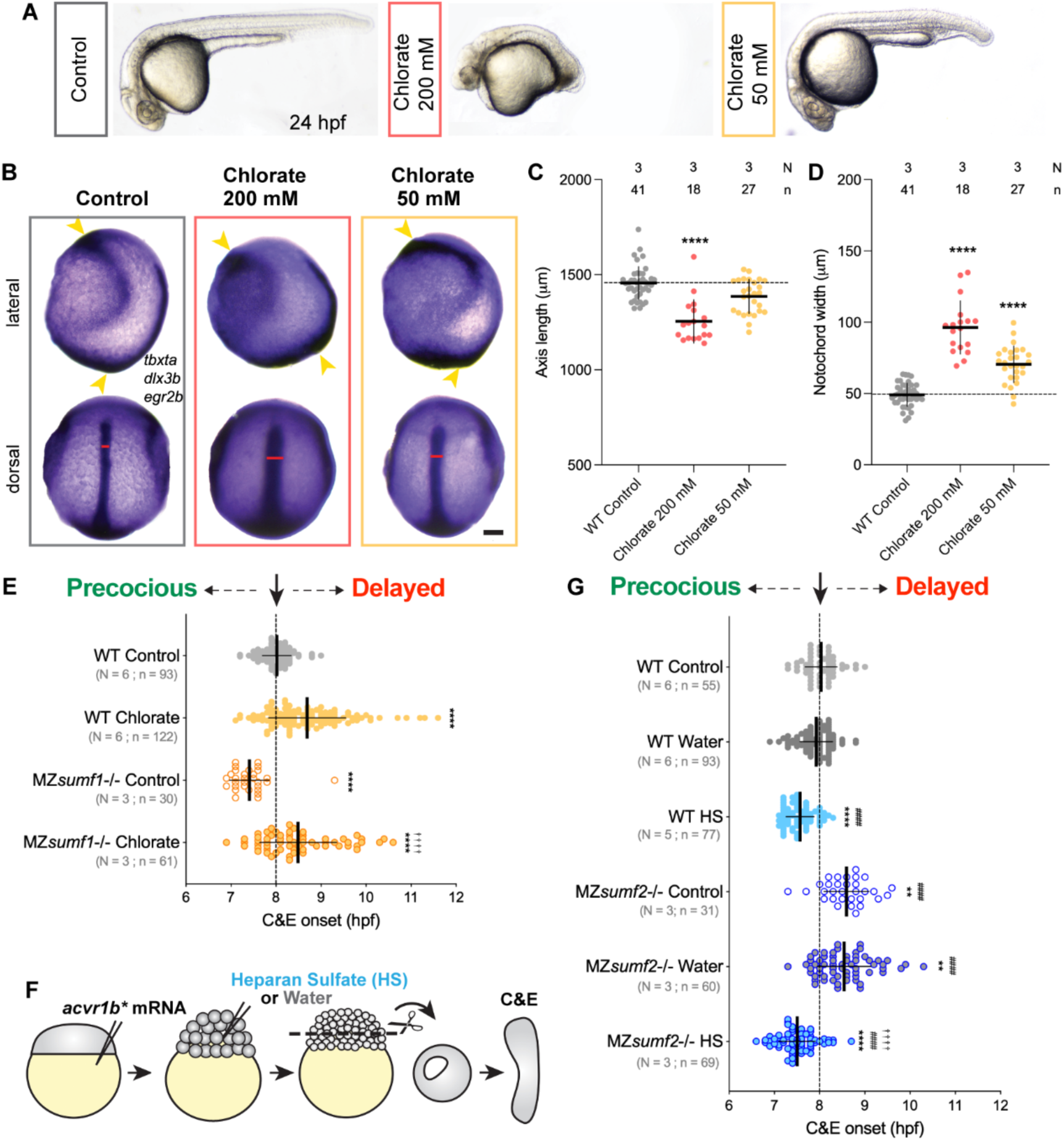
HSPGs mediate the effect of sulfatase modifying factors on C&E timing. **(A)** Representative images of 24 hpf zebrafish embryos treated at dome stage (4.33 hpf) with 200 mM or 50 mM of sodium chlorate. **(B)** Representative images of WISH for *tbxta* (mesoderm), *dlx3b* (neural border) and *egr2b* (rhombomeres 3 & 5) in tailbud stage (10 hpf) control and sodium chlorate treated embryos shown from lateral (top) and dorsal (bottom) views. Scale bar 100 μm. **(C, D)** AP axis length (C) and Notochord width (D) (as in Figure 2) of embryos depicted in (B). Each dot represents a single embryo. Means and standard deviation are indicated. **** p<0.0001 compared with WT control embryos by Kruskal–Wallis and Dunn’s multiple comparisons tests. **(E)** Onset of extension in *acvr1b\** explants of the indicated conditions. Dotted line shows typical extension onset of WT control explants around 8 hpf. Each dot represents a single explant. Means and standard deviation are indicated. N: number of independent experiments, n: number of embryos. ****^/††††^ p<0.0001 compared with WT and MZ*sumf1*-/- control groups, respectively, by Kruskal–Wallis and Dunn’s multiple comparisons tests. **(F)** Diagram of extracellular heparan sulfate (HS) or water (control) injections in 256-cell embryos prior to explantation (modified from (*36*)). **(G)** Explant extension onset as in (E). ** p>0.01; ****^/####/††††^ p<0.0001 compared with WT, WT + water, and MZ*sumf2*-/- + water groups, respectively, by Kruskal–Wallis and Dunn’s multiple comparisons tests.

**Supplementary Fig. 5.**
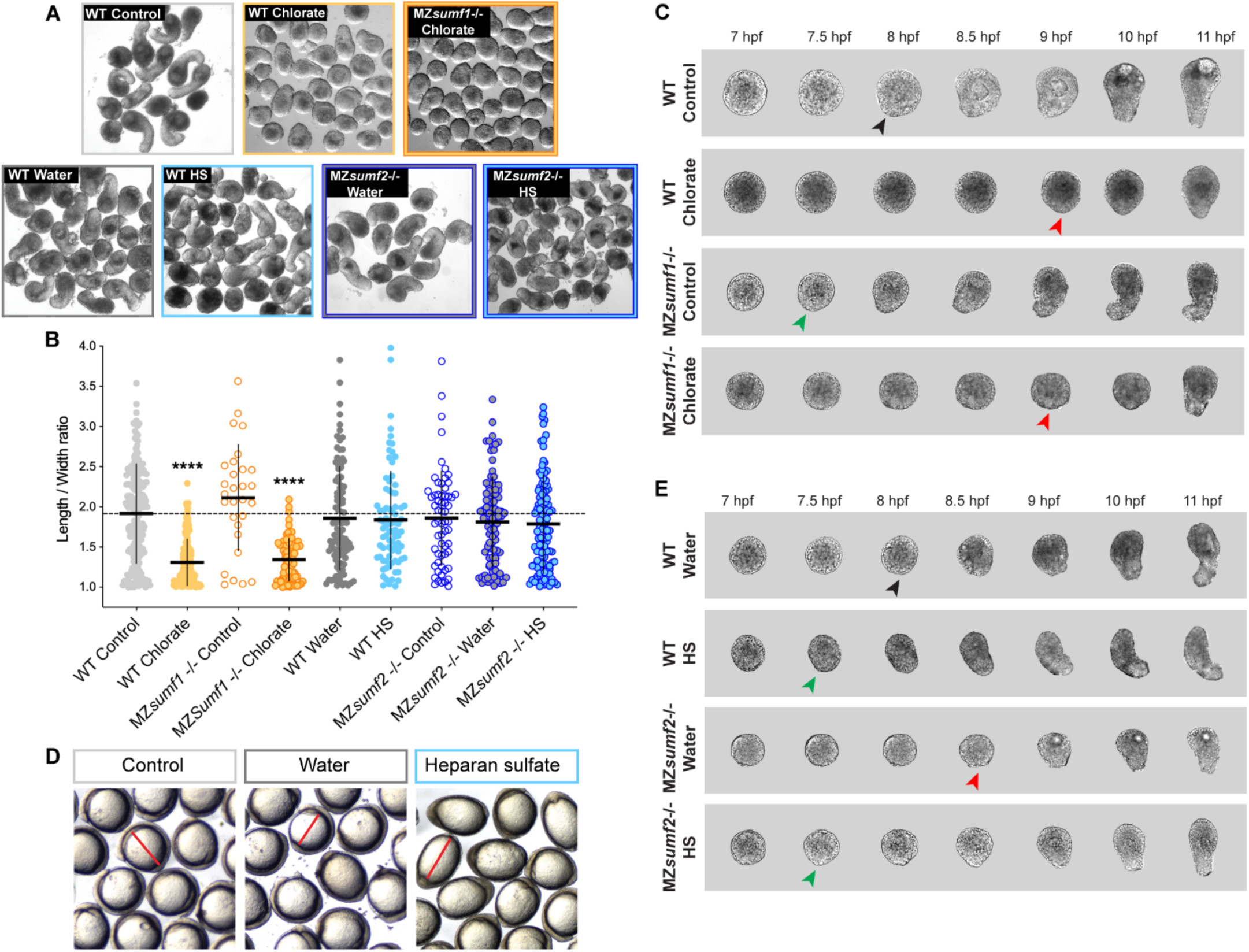
Inhibition of sulfation, but not increased heparan sulfate levels, impairs C&E *ex vivo*. **(A)** Representative images of *acvr1b\** explants from the indicated conditions. **(B)** Length/width ratios of explants shown in (A). Each dot represents a single explant. Means and standard deviation are indicated. **** p<0.0001 compared to WT controls by Kruskal–Wallis and Dunn’s multiple comparisons tests. **(C)** Representative bright-field images of *acvr1b\** explants of the indicated genotypes and treatments over time. Black, red, and green arrowheads indicate timely, precocious, and delayed onset of extension, respectively. **(D)** Heparan sulfate (HS) injected embryos display a dorsalized phenotype at the 2-somite stage (10.66 hpf), indicated by increased animal–vegetal length (red lines). **(E)** Representative bright-field images of *acvr1b\** explants of the indicated genotypes and treatments over time, as in (C).

In a complementary experiment, we increased levels of sulfated HS by injecting the extracellular space of 256-cell stage embryos with 10 ng/ml HS (or water as a control) (**Fig. 6F**). As previously reported (*92*), HS-injected embryos exhibited dorsalized phenotypes (**Supplementary Fig. 5D**), likely reflecting increased FGF signaling (*93*). Notably, while increased HS did not affect the extension of *acvr1b⁎* explants at the equivalent of 4-somite stage (**Supplementary Fig. 5A, B**), their extension onset was significantly earlier than control explants (**Fig. 6G, Supplementary Fig. 5E**). HS injection also advanced extension onset in MZ*sumf2*-/- explants (**Fig. 6G, Supplementary Fig. 5E**), indicating that increased HS levels are sufficient to trigger precocious C&E *ex vivo* and can override the delay caused by *sumf2* deficiency. Taken together, these results support a model in which *sumf2* expression at gastrulation triggers the timely onset of C&E cell movements by reducing Sulf1 activity and consequently increasing sulfation of HSPGs.

## DISCUSSION

Morphogenesis requires precise temporal coordination of cell behaviors to ensure proper tissue shape and function, however, the molecular mechanisms governing its timing remain poorly understood. In this study, we uncover a novel molecular mechanism controlling the timing of C&E morphogenesis. After establishing that transcription is required at gastrulation onset for C&E, we identified the sulfatase modifying factors *sumf1*/FGE and *sumf2*/pFGE as key temporal regulators. Our data support a model (**Fig. 7**) in which, prior to gastrulation, maternally deposited *sumf1*/FGE activates sulfatases enzymes, including the extracellular sulfatase Sulf1. At gastrulation onset, *sumf1* and *sumf2* transcript abundance inverts, and the antagonistic effect of *sumf2*/pFGE on *sumf1*/FGE reduces Sulf1 activity. Consequently, increased levels of sulfated HSPGs promote and/or permit C&E morphogenesis.

**Fig. 7.**
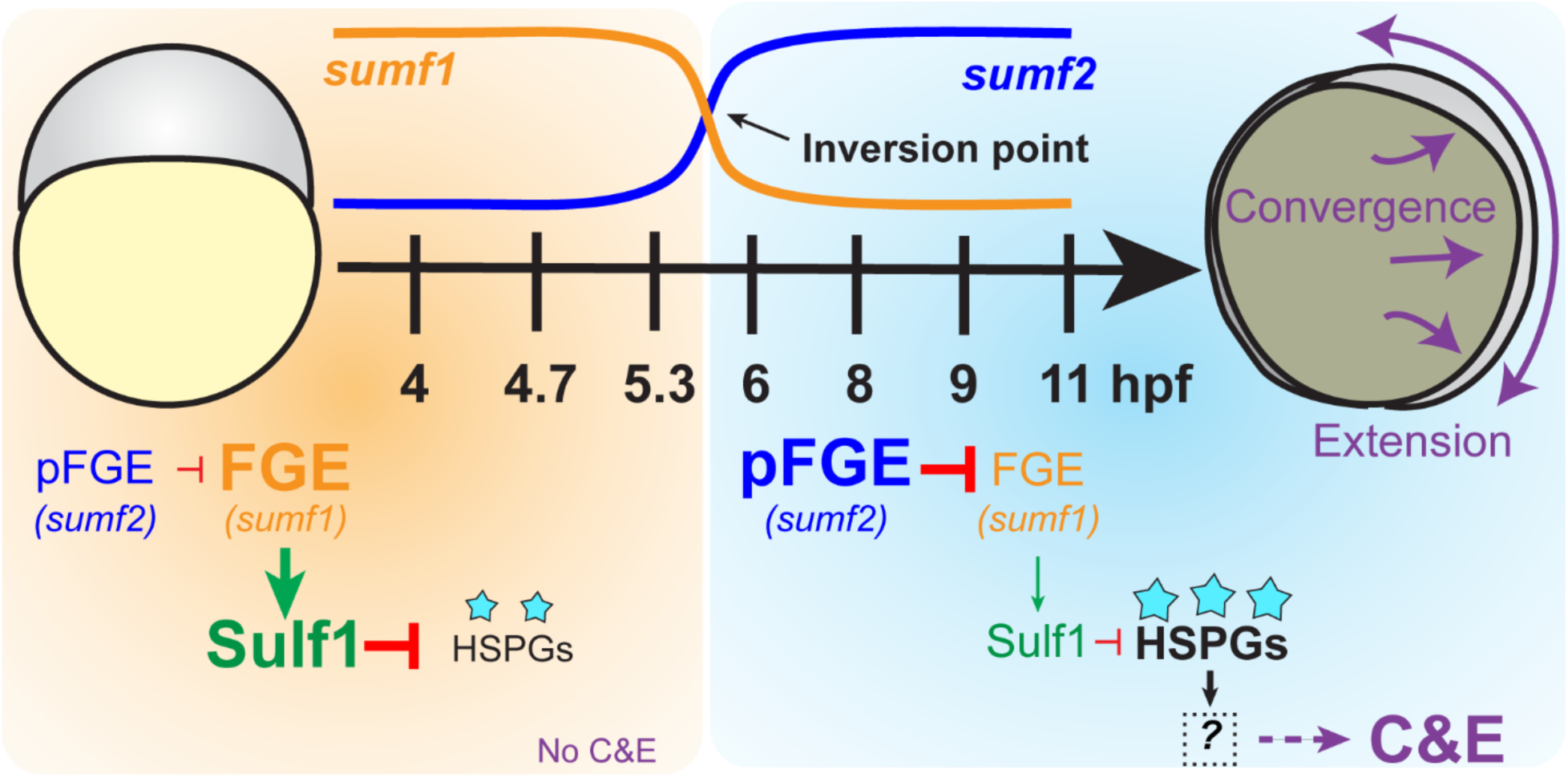
A model for how sulfatase modifying factors regulate the timing of C&E morphogenesis. Before gastrulation (orange), maternally expressed *sumf1*/FGE activates Sulf1, maintaining low levels of 6-*O*-sulfation (cyan stars) on HSPGs. At the onset of gastrulation (light blue), an inversion in *sumf1* and *sumf2* transcript abundance (inversion point) increases the antagonistic action of *sumf2*/pFGE on *sumf1*/FGE, thereby reducing Sulf1 activity. This shift leads to increased 6-*O*-sulfation of HSPGs, generating a permissive environment for C&E morphogenesis via an unknown downstream mechanism.

### Developmental roles of sulfatase modifying factors

This is (to our knowledge) the first report of a role for sulfatase modifying factors in vertebrate gastrulation. In humans, mutations in *SUMF1* cause multiple sulfatase deficiency (MSD), a rare and fatal autosomal recessive disorder characterized by lysosomal dysfunction, developmental delay, neurodegeneration, and skeletal defects including scoliosis, facial dysmorphism, and growth retardation (*55, 94*). Mouse *Sumf1*-/- models exhibit early postnatal lethality (*95*), and MZ*sumf1*-/- zebrafish recapitulate some MSD features such as cranial malformations and early growth retardation (*78*). However, these fish can survive to adulthood, suggesting the presence of alternative mechanisms of sulfatase activation in this species (*78*). Indeed, *Escherichia coli, Caenorhabditis elegans,* and *Sacchromyces cerevisiae* possess sulfatase genes but lack a Sumf1 homolog, suggesting the existence of an alternative formylglycine generating enzyme in these organisms (*56*). Whether a similar alternative enables MZ*sumf1*-/- zebrafish to survive is unknown.

The role of *sumf2*, on the other hand, had not been studied in vertebrate development. Like MZ*sumf1*-/- mutants, we found that MZ*sumf2*-/- fish can reach adulthood and some individuals exhibit scoliosis and craniofacial abnormalities (**Supplementary Fig. 2E**). This indicates that although *sumf2*/pFGE has a significant role in the timing of C&E morphogenesis, it is not ultimately essential for life in zebrafish. Whether its loss is compatible with life in other vertebrate species remains to be tested. Although Sumf1 orthologs are present in prokaryotes, Sumf2 is restricted to eukaryotes and has been lost in several metazoan clades, including arthropods (*55, 75, 94, 96*). *sumf2*/pFGE and *sumf1*/FGE amino acid sequences are highly similar, but *sumf2*/pFGE lacks formylglycine generating activity due to the absence of an active site (*55, 59, 60*). It was shown in human tissue culture that *sumf2*/pFGE reduces *sumf1*/FGE enzymatic activity (*60, 65*), which is thought to occur through direct physical binding and possible interactions with sulfatase enzymes (*59, 65*). This antagonism is consistent with our findings in zebrafish that MZ*sumf2*-/- phenotypes are exacerbated by *sumf1* OE (**Fig. 2**) and co-overexpression of both factors offsets the effects of their individual overexpression (**Fig. 3**). Further, MZ*sumf1*-/-; MZ*sumf2*-/- double mutant explants phenocopy the C&E delay observed in MZ*sumf1*-/-, indicating that *sumf1* is epistatic to *sumf2* (**Fig. 3**), consistent with a model in which *sumf2*/pFGE governs gastrulation cell movements in its capacity as a *sumf1*/FGE inhibitor.

### Sulf1 and HSPGs regulate gastrulation cell movements

Our data implicate Sulf1 as the primary sulfatase affecting C&E timing downstream of *sumf1*/FGE and *sumf2*/pFGE regulation. Sulf1 catalyzes the removal of 6-*O*-sulfation from the D0S6 and D2S6 disaccharides of HSPGs (*67*), precisely the modifications that were increased between early and late zebrafish gastrulation (**Fig. 5**). Notably, this function is also served by the extracellular sulfatases Sulf2a and Sulf2b. However, *sulf2b* OE did not cause pronounced gastrulation phenotypes, and although *sulf2a* OE induced C&E defects in both embryos and explants (**Supplementary Fig. 4**), it did not (unlike *sulf1* OE) alter the onset of explant extension. Although these sulfatases target the same substrate, evidence indicates that human Sulf1 and Sulf2 act on distinct polysaccharide substrates (*97*), which may also explain the differential effects of *sulf1* and *sulf2a/b* in zebrafish gastrulae. While the contribution of other sulfatases cannot be excluded, our finding that *sumf1* overexpression no longer affects C&E onset in the absence of *sulf1* strongly implicates Sulf1 as the key regulator of C&E timing. It is less clear that reduced Sulf1 activity is predominantly responsible for the overall increase in HS levels in later gastrulation, as the abundance and sulfation patterns of HS chains are ultimately determined by the activity of multiple enzymes, including glycosyltransferases, heparinases, sulfatases and sulfotransferases (*98*). Interestingly, *xylosyltransferase I* (*xylt1*), whose activity initiates HS biosynthesis (*97*), was also among our candidate ‘trigger’ genes upregulated at gastrulation onset (**Fig. 1**), making it an especially compelling candidate for further investigation.

HSPGs and the sulfatases that modify them have long been known to regulate gastrulation morphogenesis. For example, elimination of HS chains or loss of HSPG core proteins glypicans and syndecans caused gastrulation defects and/or shortened axes in *Xenopus*, zebrafish, and sea urchin gastrulae (*84, 89, 90, 99–103*). Although our study does not address the HSPG core protein(s) responsible for the observed effects on C&E timing, glypicans and syndecans with established roles in C&E (like *gpc4/kny* (*101*)) are good candidates. Notably, our manipulations affecting HSPG sulfation tended to produce milder phenotypes that loss of core proteins like Gpc4, consistent with sulfation as a modifier of their activity. Indeed, manipulations that decrease sulfation levels – including overexpression of *sulf1* or *sumf1* mRNA and injection of purified sulfatases into the blastocoel - also disrupted gastrulation in *Xenopus* and sea urchin (*64, 80–82*). Interestingly, loss of sulfatases also caused gastrulation defects (*82, 104, 105*), suggesting that a correct balance of sulfatase activity is required for proper C&E. Our data provide a possible explanation for this: altered timing of C&E movements. We found that *sulf1* overexpression both disrupts and delays C&E, but also that both precocious and delayed C&E onset ultimately cause C&E defects (i.e. shorter and wider embryonic axes) in intact embryos. This raises the possibility that previously reported C&E defects upon gain or (especially) loss of sulfatase function could be the result of altered timing (discussed further below).

*sulf1* deficiency, however, not only alters the timing of C&E but also prevents full explant extension (**Fig. 4, Supplementary Fig. 4**). This discrepancy in phenotypic severity between Sulf1 and its key regulator (*sumf1*/FGE) may reflect partially independent regulation of sulfatases during zebrafish gastrulation. As discussed above, C&E defects resulting from altered sulfatase activity are relatively mild in zebrafish, and the mutant embryos examined here can ultimately survive to become fertile adults. This suggests the existence of compensatory mechanisms that support continued primary body axis elongation after gastrulation is complete.

### Temporal dynamics of gastrulation cell movements

Although it is intuitive that delayed C&E ultimately manifests as a C&E defect *in vivo*, it is less clear how increased and/or precocious convergence movements (as seen in MZ*sulf1*-/-, MZ*sumf1*-/-, and *sumf2* OE embryos) lead to similar phenotypes (**Fig. 3 and Fig. 4**). We speculate that all gastrulation cell movements – epiboly, internalization, and C&E – must be coordinated in both space and time to properly shape the nascent body axes. If C&E movements are accelerated, they become out of sync with other concurrent cell movements, disrupting axis extension. This would explain why morphological C&E defects are detected in MZ*sumf1*-/- mutants but not explants (**Fig. 2 and Supplementary Fig. 3**), in which C&E occur in the absence of epiboly and internalization. A similar phenomenon was reported in *Drosophila* gastrulae, in which both slowed and accelerated germ band extension desynchronized morphogenesis between the three germ layers (*106, 107*).

It is not yet clear whether sulfatase activity alters the onset of cell movements, their pace, or both. For example, either accelerated cell movements or their precocious onset could manifest as “early onset” of morphological extension within our explants. Indeed, *Drosophila* embryos with loss- and gain-of function mutations in Serotonin signaling components reportedly exhibit slowed and accelerated cell movements driving germ band extension, respectively, without a change in their time of onset (*106–108*). However, our *in vivo* cell-tracking analysis revealed apparent changes in both the onset and speed of convergence movements in zebrafish gastrulae with altered *sumf1, sumf2*, and *sulf1* levels, suggesting that multiple aspects of ‘timing’ are affected (**Fig. 3 and Fig. 4**). The precise changes in the timing of cell behaviors upon altered sulfatase activity, and their contributions to our observed phenotypes, are areas of interest for future study.

### How do HSPGs govern morphogenetic timing?

Cell surface and extracellular HSPGs play key roles in multiple morphogen signaling pathways. For example, HSPGs modulate ligand diffusion and availability (*67, 81, 109, 110*) and receptor binding (*67, 81, 111*) of morphogens with known roles in C&E, including FGF, Wnt, BMP, and Nodal. HSPGs also regulate PCP signaling through binding of non-canonical Wnt ligands (*99, 101, 111*) and membrane localization of Dishevelled (*81, 99, 100*), which is required for PCP activity (*112, 113*). The structural features of HS chains, particularly their sulfation patterns, are critical for these functions as they determine ligand-binding affinity and selectivity (*98, 114, 115*). Notably, the precise role of HSPG sulfation in signaling is context-dependent and changes throughout development. For example, reduced sulfation inhibits and enhances signaling by Wnt8 and Wnt11, respectively (*81, 82, 116, 117*), and HSPGs from different developmental stages have different capacities to bind FGF ligands (*118, 119*). We speculate that sulfatase modifiers may regulate C&E timing by controlling HS sulfation, thereby modulating the degree and dynamics of HSPG-mediated signaling. Indeed, we observed an increase in the levels of sulfated HSPGs (Sulf1 substrates) during late gastrulation (**Fig. 5**) and demonstrated that both decreased and increased levels of sulfated HS were sufficient to alter C&E timing (**Fig. 6**). Importantly, the phenotypes associated with *sumf1/2* and *sulf1* perturbations reflect disrupted morphogenesis in the absence of obviously altered cell fate specification, as embryos still give rise to derivatives of all three germ layers, consistent with observations in *Xenopus* and sea urchin gastrulation (*80, 84*). However, we cannot exclude the possibility that the alterations in C&E timing are secondary to subtle patterning defects or changes in the timing of cell fate choices. Whether this putative role in morphogen signaling underlies the effect of sulfatase modifiers on C&E timing will be an exciting topic for future studies.

## METHODS

### Zebrafish

Adult zebrafish were maintained following established protocols (*120*) in compliance with Baylor College of Medicine Institutional Animal Care and Use Committees. Embryos were obtained through natural mating, and staging was based on established morphology (*121*). Fish were chosen from their home tank to be crossed at random, and embryos were randomly chosen from the clutch for injection and inclusion in experiments. WT and mutant embryos were collected in parallel in a period not longer than 10–15 min to minimize developmental differences and raised in egg water at 28.5°C under identical conditions. Experiments on WT embryos were conducted in either the AB or TU background, depending on the background of the mutant line used. The mutant lines used were *sum^la015919Tg^*(ZFIN ID: ZDBALT-120806-11568), *sulf1^sjr9^* (Kaur et al, submitted), *sumf2^bcm126^* (this study, described further below) and *oep^tz257^* (*101*)*. oep-/-* embryos were rescued to viability by injection of 50 pg of *oep* mRNA (*122*) and raised to adulthood, then intercrossed to generate MZ*oep-/-* embryos for explantation. MZ*sumf1*-/-; MZ*sumf2*-/- double mutants were generated by crossing single homozygous mutants to obtain double heterozygotes, which were subsequently incrossed.

### *sumf2^bcm126^* generation and genotyping

*sumf2* zebrafish mutants were generated using the CRISPR–Cas9 system. A single guide (sg)RNA targeting exon 2 of *sumf2* (5′-GGATGGAGAATCGCCAACAC-3′) was designed using CRISPRscan (*123*). sgRNAs were transcribed using T7 RNA polymerase (NEB, M0251S) from DNA templates generated with the forward primer (Fw): gaaattaatacgactcactataGGATGGAGAATCGCCAACACgttttagagctagaaatagc, and the reverse primer (Rv): aaaagcaccgactcggtgccactttttcaagttgataacggactagccttattttaacttgctatttctagctctaaaac. 1 μl of gRNA was pre-incubated for 10 min at 37 °C with Cas9 protein (NEB, M0646M) and 300mM KCl (as described by (*124*)). AB WT Embryos were injected at the single-cell stage with 1 nl of the gRNA–Cas9 complex and cutting efficiency was assessed using a T7 endonuclease I assay (*125*). A PCR fragment encompassing the target sequence was amplified from genomic DNA (gDNA) of individual embryos (using the following primers: Fw: AGATGGTGTTTATTCCTGGTGG, Rv: TCCTCTGATACAAAATCCTGGAA) and incubated with T7 endonuclease I (NEB #M0302L) in NEB 10× Buffer 2. Injected F0 embryos were raised and outcrossed to AB WT fish. F1 progeny were genotyped by Sanger sequencing to identify transmitted mutations. A 7 bp insertion in exon 2 of *sumf2* (sumf2*^bcm126^)*, resulting in an early stop codon, was selected (**Supplementary Fig. 2A**). This mutation introduced an XcmI restriction site, enabling genotyping by digestion of the PCR-amplified target fragment with XcmI (NEB, R0533S) (**Supplementary Fig. 2B**). Heterozygous F1 carriers were incrossed to produce the F2 generation. Homozygotes F2 fish were raised and incrossed to obtained MZ*sumf2*-/- embryos for experiments.

### RT-qPCR

RNA was isolated from 50 pooled WT and MZ*sumf2*-/- 24 hpf larvae from three independent clutches. Total RNA was extracted using TRIzol reagent (Thermo Fisher Scientific, 15596018), purified by sodium acetate precipitation, eluted in 10 mM Tris pH7.5 and treated with Turbo DNAase I (Invitrogen, AM2238) at 37°C for 30 min. For each sample, 1μg of the RNA was reverse-transcribed using iScript Reverse Transcription Supermix for RT-qPCR (Biorad, #1708840). *sumf2* specific primers were designed to span exon-exon junctions [Exon 1-2 (pair 1) FW: CACAGTGTCTTGTGCAGCAG, RV: AGTTGGAGTTGGTGACAGGA; Exon 3-5 (pair 2) FW: GGCTGAAACATTTGGCTGGA, RV: GGCATCATTCCAGCTGACCT]. *rpl13a-1* was used as a housekeeping gene [FW: TCTGGAGGACTGTAAGAGGTATGC, RV: AGACGCACAATCTTGAGAGCAG]. qPCR was performed in technical triplicates using SsoAdvanced Universal SYBR Green Supermix (Biorad, #1725270). Relative expression levels were calculated using the 2^−ΔΔCt^ method.

### Preparation and microinjection of mRNA

All mRNAs were transcribed using the SP6 mMessage mMachine Kit (Fisher Scientific, AM1340) and purified using Bio -Rad Microbiospin columns (Bio-Rad, 7326250). Single-celled embryos were placed in agarose molds (Adaptive Science Tools, I-34) and injected with 0.5-1 nl volumes using pulled glass needles (Fisher Scientific, 50-821-984). Doses of mRNA per embryo were as follows: 0.5 pg *acvr1b\** (*69*), 10 pg *ndr2* (*126*), 50-100 pg s*umf1*, 100 pg *sumf2*, 500 pg *sfGFP,* 500 pg *sulf2b,* 500 pg *sulf2a,* 500pg *gnsb,* 50-500 pg *sulf1,* 500 pg *sgsh,* 500 pg *arsb,* 500 pg *gnsa,* 500 pg *ids,* 500 pg *galns,* 500 pg *arsa,* 500 pg *sts,* 100 pg *mem-GFP* and 100 pg *H2B-scarlet.* Templates for *sumf1*, *sumf2*, and all sulfatases were generated by Gibson cloning (*127*) each of their open reading frames (synthesized by Twist Biosciences) into a PJS2 vector linearized with EcoRI.

### Whole mount *in situ* hybridization (WISH)

*tbxta (brachyury, t)*, *dlx3b* and *egr2b (krox20)* antisense riboprobes were transcribed using NEB T7 RNA polymerase (NEB, M0251s) and labeled with digoxigenin NTPs (Sigma/Millipore, 11277073910) NTPs (Sigma/Millipore, 11685619910). WISH was performed according to (*128*). Embryos were fixed overnight in at 4°C 4% PFA in PBS, rinsed in PBS + 10% tween-20 (PBT), and dehydrated into methanol. Following rehydration into PBT, embryos were pre-incubated for at least two hours in hybridization buffer with 50% formamide (in 0.75 M sodium chloride, 75 mM sodium citrate, 0.1% tween 20, 50 mg/mL heparin (Sigma), and 200 mg/mL tRNA) at 70°C, and hybridized overnight at 70°C with antisense probes (1-5 ng/mL) in hybridization buffer. Samples were washed gradually into 2X SSC buffer (0.3 M sodium chloride, 30 mM sodium citrate), and then gradually from SSC to PBT. Samples were blocked at room temperature for several hours in PBT with 2% goat serum and 2 mg/mL bovine serum albumin (BSA), then incubated overnight at 4°C with anti-DIG antibody (Roche #11093274910) at 1:5000 in block. After extensive washes in PBT, embryos were rinsed in staining buffer (PBT +100 mM Tris pH 9.5, 50 mM MgCl_2_, and 100 mM NaCl) and developed in BM Purple AP substrate (Roche) until the desired staining intensity was achieved. Staining was stopped with 10 mM EDTA in PBT before imaging.

### Blastoderm explants

Blastoderm explants were prepared as described by (*129*). Briefly, uninjected, *acvr1b*-,* or *ndr2*-injected embryos were dechorionated at the 256-cell stage using pronase (Roche; 1 ml of 20 mg/ml stock in 15 ml 3× Danieau’s solution). At the 512-cell stage, approximately 1/3 of the most animal blastoderm cells were excised using Dumont #55 forceps (Fisher Scientific, NC9791564) on an agarose-coated plate containing 3× Danieau’s solution. Explants were allowed to heal briefly before being transferred into agarose-coated 6-well plates containing explant medium [Dulbecco’s modified eagle medium with nutrient mixture F-12 (Gibco 11330032) containing 2.5 mM L-glutamine, 15 mM HEPES, 3% newborn calf serum (Invitrogen 26010–066), 50 units/mL penicillin, and 50 mg/mL streptomycin (10,000 U/mL pen-strep at 1:200, Gibco 15140163)]. Explants were incubated at 28.5 °C until sibling embryos from the same genetic background reached the desired developmental stage.

### Alcian Blue staining

Embryos were fixed in PFA 4% overnight, extensively washed in PBS, and incubated in Alcian Blue staining solution pH 1 (0.2% Alcian Blue 8 GX (Sigma # A5268), 50% ethanol, ∼0.1N HCl) for 48 h at room temperature, protected from light. Samples were then gradually rehydrated into PBS, post-fixed in 4% PFA, washed with PBS, and rinsed in 2% KOH. Embryos were cleared through a graded series of glycerol in 2% KOH (20%, 40%, 60%) and stored in 80% glycerol in 2% KOH for imaging and long-term storage.

### Transcription inhibitor treatment

1 μM Triptolide (Sigma # T3652) or an equivalent volume of DMSO was added to the media of blastoderm explants in agarose-coated 6-well plates at 50% (4.7 hpf), shield (6 hpf), 70% (7.5 hpf) or 80% (8.5 hpf). 0.5 μM Flavopiridol (Selleck, S1230) or an equivalent volume of DMSO was added to the media of blastoderm explants at 50% (4.7 hpf) and washed-out twice with 0.3x Danieau solution, before incubation with fresh explant medium.

### Sodium Chlorate treatment

Dome stage (4.33 hpf) embryos or blastoderm explants were treated with 200 or 50 mM of Sodium Chlorate (VWR # 7775-09-9) in egg water or explant medium.

### Heparan Sulfate injections

Uninjected or *acvr1b\**-injected embryos were dechorionated at the 64-cell stage using pronase (Roche; 1 mL of 20 mg/mL stock in 15 mL 3× Danieau’s solution). Embryos were transferred to agarose-coated plates containing cubical depressions (Adaptive Science Tools, PT-1) filled with 0.3× Danieau’s solution. At the 256-cell stage, embryos were injected with 2 nL of 5 ng/mL heparan sulfate (Sigma-Aldrich, H7640) or nuclease-free water into the extracellular space.

### Microscopy

Live embryos injected with *H2B-mScarlet* and *mem-GFP* mRNA were manually dechorionated and mounted in 0.3-0.35% low-melt agarose (Thermo Fisher Scientific, 16520100) in glass-bottomed 35 mm Petri dishes (Fisher Scientific, FB0875711YZ) for imaging using a Nikon ECLIPSE Ti2 inverted confocal microscope equipped with a Yokogawa W1 spinning disk unit, PFS4 camera, and 405/488/561 nm lasers (emission filters: 455/50, 525/36, 605/52). Temperature was maintained at 28.5°C during imaging using a Tokai Hit STX stage top Incubator. For live time-lapse series, 100 μm z-stacks with a 2 μm step were collected every 5 minutes for 5 hours using a Plan Apo Lambda 20x dry objective lens. Live blastoderm explants were mounted in rounded chambers made in 1% low-melt agarose in glass-bottomed dishes containing explants medium and imaged in the same scope. For live time-lapses, 14 μm z-stacks were obtained with a 2 μm step every 10 minutes for 6 hours using a Plan Apo Lambda 10x dry objective lens. Images of WISH and alcian blue-stained embryos and live embryos, larvae and explants were taken with a Nikon Fi3 color camera on a Nikon SMZ745T stereoscope.

### Image analysis

ImageJ/Fiji was used to visualize and measure all microscopy data sets.

#### Morphometric analyses

During analysis, researchers were kept unaware of the conditions of all image data using the blind_renamer Perl script (https://github.com/jimsalterjrs/blindanalysis) (blindanalysis: v.1.0.) prior to analysis. To measure the length/width ratios of explants, the length of a segmented line drawn along the midline of each explant (accounting for curvature) was divided by the length of a perpendicular line spanning the maximal width of the explant. Tailbud morphometries were performed in whole-mount embryos staged using *egr2b* expression at the future mid-hindbrain boundary. Notochord width was quantified in dorsal-view images as the mediolateral extent of the *tbxta* expression domain at the midline. Anteroposterior (AP) axis length was measured in lateral-view images from the anterior to posterior boundaries of the *dlx3b* expression domain.

#### Explant onset of extension

The onset of explant extension was assessed in a blinded manner as the time point when a visible tip first emerged from an initially rounded explant. Explant were staged according to sibling intact embryos from the same genetic background cultured in the same plate.

#### Cell tracking analysis

Automated nuclear tracking was performed using the ImageJ TrackMate7 plugin (*130*) in the dorsal hemisphere (encompassing dorsal and lateral cells) of zebrafish gastrulae injected with *H2B-scarlet* mRNA. TrackMate generated color-coded trajectories and measurements of mediolateral (ML) displacement. Embryos were staged relative to formation of the second somite. To minimize noise from reduced convergence movements near the dorsal midline, cells within 100 μm of the midline were excluded from the analysis. Average displacement in the X (ML) dimension was calculated per time frame, smoothed using a sliding window of four time points, and plotted over time using GraphPad Prism 10.

### GAG isolation and disaccharide analysis

Whole zebrafish gastrulae (5.3 or 8.5 hpf, 100 embryos/sample) were homogenized and lysed in 0.5% CHAPS lysis buffer (50 mM HEPES, 120 mM NaCl, 2 mM EDTA, pH 7.4) containing a protease inhibitor cocktail (Roche). 50 µL of cell lysate was set aside for protein quantification via BCA assay. Homogenates were diluted 1:10 in a wash buffer (50 mM sodium acetate, 200 mM NaCl, 0.1% Triton X-100, pH 6.0) and incubated with Pronase (0.4 mg/ml, Sigma) overnight at 37 °C with mild agitation. The product was centrifuged (4,000 xg, 20 minutes) then passed through a DEAE-Sephacel (Cytiva) column equilibrated in 50 mM sodium acetate buffer, pH 6.0, containing 200 mM NaCl, and desalted using a PD-10 desalting column (Cytiva). The purified GAGs were subsequently enzymatically depolymerized with 2 mU each of heparin lyases I-III (IBEX) and differentially mass labeled by reductive amination with aniline, as described (*87*). Tagged HS disaccharides were analyzed by HILIC-Q-TOF-MS on a Waters Synapt XS Q-TOF mass spectrometer, as previously described (*87*). Samples were quantified using isotopically labeled internal disaccharide standards and normalized to total protein, as measured by BCA.

### Statistical analysis

Number of embryos (n) and independent experimental replicates (N) for animal studies are stated in graphs. All experiments were performed at least in triplicates. GraphPad Prism 10 software was used to perform statistical analyses and generate graphs for all data analyzed. Datasets were tested for normality prior to analysis and statistical tests were chosen accordingly. The statistical tests used for each data set are noted in figure legends.

## ACKNOWLEDGEMENTS

We thank Dr. Lila Solnica-Krezel for sharing plasmids and WISH probes, the BCM Center for Comparative Medicine for taking excellent care of our fish, and the Zebrafish International Resource Center for preserving and distributing fish lines used here and by countless members of the community. Thanks also to all members of the Williams lab for their help and feedback on this project, and Drs. Maria Cecilia Cirio and Lance Davidson for their thoughtful comments on the manuscript.

## AUTHOR CONTRIBUTIONS

A.S.C. and M.K.W. conceived of the project. A.S.C. and M.K.W. performed zebrafish experiments, R.J.W. and A.B. performed HS disaccharide profiling, and R.M.J. and G.K.B. generated the *sulf1* deletion line. S.G. and C.C. performed bioinformatic analysis. A.S.C and M.K.W. wrote the original manuscript. All authors reviewed the manuscript.

